# Free Energy Landscape and Rate Estimation of the Aromatic Ring Flips in Basic Pancreatic Trypsin Inhibitor Using Metadynamics

**DOI:** 10.1101/2021.01.07.425261

**Authors:** Mandar Kulkarni, Pär Söderhjelm

## Abstract

Aromatic side-chains (phenylalanine and tyrosine) of a protein flip by 180° around the *C_β_ − C_γ_* axis (*χ*_2_ dihedral of side-chain) producing two symmetry-equivalent states. The ring-flip dynamics act as an NMR probe to understand local conformational fluctuations. Ring-flips are categorized as slow (ms onwards) or fast (ns to near ms) based on timescales accessible to NMR experiments. In this study, we investigated the ability of the infrequent metadynamics approach to discriminate between slow and fast ring-flips for eight individual aromatic side-chains (F4, Y10, Y21, F22, Y23, F33, Y35, F45) of basic pancreatic trypsin inhibitor (BPTI). Well-tempered metadynamics simulations were performed to observe ring-flipping free energy surfaces for all eight aromatic residues. The results indicate that *χ*_2_ as a standalone collective variable (CV) is not sufficient to classify fast and slow ring-flips. Most of the residues needed *χ*_1_ (*N − C_α_*) as a complementary CV, indicating the importance of librational motions in ring-flips. Multiple pathways and mechanisms were observed for residues F4, Y10, and F22. Recrossing events are observed for residues F22 and F33, indicating a possible role of friction effects in the ring-flipping. The results demonstrate the successful application of the metadynamics based approach to estimate ring-flip rates of aromatic residues in BPTI and identify certain limitations of the approach.

## INTRODUCTION

The fluctuations of protein residues in its native state produce an ensemble of interconvertible substates.^1^ These dynamic protein fluctuations occur over a wide range of timescales. For example, the ring-flip motions (180° rotation around C_β_-C_γ_ axis) of aromatic side-chains in globular proteins occur from nanoseconds^2^ to seconds^3^. The presence of different timescales for aromatic ring-flipping in proteins was initially observed in seminal NMR studies by Wüthrich and co-workers^4–5^ while studying the internal dynamics of basic pancreatic trypsin inhibitor (BPTI). The value of the side-chain torsion angle *χ*_2_ (C_α_-C_β_-C_γ_-C_δ1_ / *CA-CB-CG-CD1* in figure 1) acts as an indicator of aromatic ring-flipping. The ring-flipping process occasionally involves librational motions relative to backbone atoms, identified using the *χ*_1_ (N-C_α_-C_β_-C_γ_ / *N-CA-CB-CG*) torsion. The aromatic side-chains are often utilized as internal NMR probes to understand the protein interior dynamics surrounding these side-chains^6–9^.

**Figure 1.**
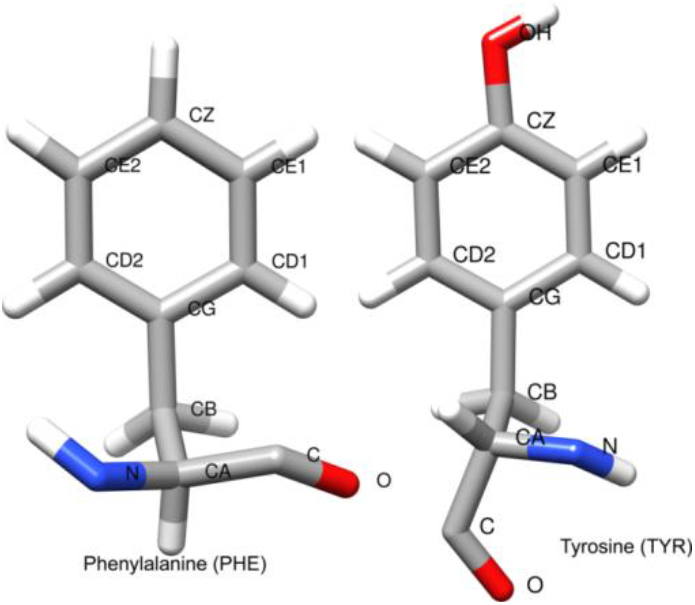
Atom names of tyrosine and phenylalanine indicating torsion *χ*_2_ (*CA-CB-CG-CD1*) and *χ*_1_ (*N-CA-CB-CG*).

The ring-flips are not directly associated with any biological functions^10^ but rely on transient packing defects around the aromatic ring, which might be relevant to small-molecule binding or internal protein hydration. The ring flipping event in the protein interior is associated with breathing motion, a transient cavity expansion exposing the protein interior. Such breathing motions provide a transient formation of the necessary volume (*activation volume*^9, 11^) required for the aromatic ring flip. This is a well-established model often referred to as the “cavity model”^12^. The earlier ring-flipping studies proposed a “diffusion model”^13–14^ based on Kramer’s theory assuming ring-flipping is a diffusion-limited process originating from the frictional effects exerted by surrounding atoms.

In this study, we investigated four tyrosine (Y10, Y21, Y23, and Y35) and four phenylalanine (F4, F22, F33, and F45) residues of BPTI. These aromatic side-chains contribute significantly to the stability of BPTI and mutations^15–17^ at these sites with alanine or leucine destabilized protein at room temperature. The NMR investigation of the globular protein BPTI by Wagner et al.^5^ was the first study to distinguish slow and fast flipping rings. In general, the aromatic ring flips are classified^5, 18^ as slow flips (k_flip_ ≤ 10^3^ s^−1^ i. e. t_flip_ ≥ 10^−3^ s) and fast flips (k_flip_ > 10^3^ s^−1^ i. e. t_fiip_ < 10^−3^ s) based on the flipping time (t_flip_) accessible to NMR methods. Experimentally, only slow ring flips can be studied using ^13^C CPMG relaxation experiments^3, 11^ by observing signal splitting of symmetrically located δ or ε nuclei on the ring^4^ (CD1, CD2, CE1, CE2 positions in figure 1). Experimental NMR studies of BPTI using ^13^C relaxation dispersion^3^ observed that the slow flipping rings Y23, Y35, and F45 in BPTI exhibit an activation enthalpy around 20 kcal/mol. The abundance of experimental observations related to all eight aromatic residues in BPTI makes it a suitable model system to understand the thermodynamics and kinetics of the ring-flipping mechanism.

Presently, it is possible to perform molecular dynamics (MD) simulations up to a few microseconds within a reasonable time due to advances in computing power and GPU based molecular dynamics codes. In a 1-ms-long BPTI (Amber99SB-I protein force field and TIP4P-Ew water model) simulation by D.E. Shaw Research (DESRES), ring-flipping was observed for seven out of eight aromatic side-chains.^19^ The sparsity of ring-flip events in DESRES simulation suggests that the ring-flipping of aromatic side-chains in BPTI is a rare event. The sampling problem of rare events is generally resolved using enhanced sampling techniques such as metadynamics^20–21^. In this study, well-tempered metadynamics^20^ (WTMetaD) simulations are performed to understand the free energy landscape of ring-flipping for all eight aromatic side-chains of BPTI. The successful application of metadynamics provides thermodynamic and structural details of the stable states. However, it is not trivial to evaluate the overall kinetics of the process using such a nonequilibrium approach. Acceleration of rare event by the application of an external biasing potential corrupts the actual dynamics of the process. Recently, the infrequent metadynamics^22^ method (InMetaD) was proposed to recover accurate transition times from slowly or *infrequently* biased metadynamics simulations. We performed InMetaD^22^ simulations to obtain the first-passage times for transitions between symmetry-equivalent flipped configurations. Analysis of the simulations allows us to propose the ring-flipping mechanism, identify intermediates, and calculated ring-flip rates.

## METHODOLOGY

### Simulation setup

The initial coordinates for the basic pancreatic trypsin inhibitor (BPTI) were obtained from PDB ID 5PTI, and deuterium atoms and water molecules were removed. The ionizable residues were modeled based on their state at pH 7. The system was built using the tleap module of *AmberTools18* and then converted to *GROMACS* format using *ACPYPE* python script^23^. The protein molecule was modeled using the AMBER ff14SB^24^ force field. The protein was first placed in the octahedron box and was solvated by 4302 four-point TIP4P-Ew^25^ water molecules. Then, six Cl^−^ ions were added to neutralize the system. The Joung-Cheatham parameters^26^ for chloride ions were used. The distance between walls and protein was 10 Å. All simulations were performed with a time step of 2 fs, and all covalent bonds were constrained using the LINCS algorithm. A cut-off distance of 10 Å was used for short-range Coulomb and Lennard-Jones interactions. A long-range dispersion correction to the energy and pressure was applied. The long-range electrostatic interactions were treated with PME and Fourier grid spacing of 1.2 Å with fourth-order spline interpolation. Periodic boundary conditions were applied, and the system was equilibrated first using the isothermal-isobaric (*NPT*) ensemble for 1.2 ns. During equilibration, the Berendsen^27^ thermostat and barostat were used to maintain 300 K temperature and 1 bar pressure. A restraining force of 25 kcal mol^−1^ Å^−2^ was used on heavy protein atoms during this stage. Then the structure was minimized using the steepest-descent method. The restraining force was reduced to 5 kcal mol^−1^ Å^−2^ and *NPT* equilibration was performed for 1 ns, followed by system minimization. After this stage, a 10 ns *NPT* equilibration run was performed using the Parrinello-Rahman barostat^28^ (coupling constant 1.0 ps) and Bussi-Donadio-Parrinello’s V-rescale^29^ thermostat (coupling constant 0.5 ps). The compressibility was set to 4.5 x 10^−5^ bar^−1^. The structure was minimized, and 100 ns of unbiased simulation was performed (*NVT* ensemble). A preliminary analysis was performed on this simulation to understand local fluctuations in BPTI. No aromatic ring-flipping was observed during this simulation. The last structure of this simulation was utilized further for metadynamics simulation as well as for all InMetaD simulations. All metadynamics simulations were performed in the *NVT* ensemble (T=300 K) using *GROMACS* 20 1 8.3^30–31^ and *PLUMED* 2.5^32–34^.

We have also tested the effect of the force field on ring-flipping for residue F22 using (i) CHARMM36m and charmm TIP3P water model (referred to as c36m) and (ii) CHARMM36 and standard TIP3P water model (referred to as c36). In c36m and c36 simulations, all covalent bonds involving hydrogen were constrained using LINCS. The long-range dispersion correction to energy and pressure was absent for CHARMM c36 and c36m simulations. For CHARMM simulations, the PME method was used to calculate long-range electrostatic interactions with a 12 Å cut-off. The Lennard-Jones interactions were smoothly switched to zero between 10 Å and 12 Å.

### Metadynamics Simulations

The experimentally fast flipping residues exhibit microsecond to near milliseconds timescales^3^, which are still difficult to access by standard molecular dynamics simulations. We have performed WTMetaD simulations to understand the aromatic ring-flipping mechanism and to observe possible intermediate states along the flipping pathway. The details of the metadynamics simulations are reported in Table 1. A cartesian coordinates (*R^N^*) based high dimensional free energy landscape of the system of interest is often studied using low dimensional or coarse-grained descriptors of the coordinate space, *s*(*R^N^*), called collective variables (CVs). The selection of CVs is critical to correctly capture the underlying free energy landscape and all possible metastable states. In the present study, the *χ*_2_ dihedral angle was the obvious choice to enhance ring-flipping. In addition, most WTMetaD simulations needed the *χ*_1_ torsion to observe convergence as librational motions relative to the backbone are needed for ring flipping.

**Table 1.** List of CVs and details of metadynamics parameters; hill width (*σ*), hill height (w, kJ.mol^−1^), bias factor (*γ*), frequency of hill addition (*v*, ps). For WTMetaD, *v* is 1 ps.

| residue | CV | $\sigma$ | WTMetaD | | InMetaD | | |
| --- | --- | --- | --- | --- | --- | --- | --- |
| | | | $\gamma$ | $w$ | $\gamma$ | $w$ | $v^a$ |
| F4 | $\chi_2, \chi_1$ | 0.07, 0.07 | 12 | 1.2 | 12 | 0.6 | 12 |
| Y10 | $\chi_2, \chi_1$ | 0.07,0.07 | 12 | 1.2 | 12 | 0.3 | 10 |
| Y21 | $\chi_2$ | 0.07 | 10 | 0.6 | 10 | 0.6 | 18 |
| F22 | $\chi_2, \chi_1$ | 0.07,0.07 | 12 | 1.2 | 8 | 1.2 | 12 |
| F22 (c36)<sup>a</sup> | $\chi_2, \chi_1$ | 0.07,0.07 | 12 | 1.2 | 8 | 0.3 | 12 |
| F22 (c36m)<sup>b</sup> | $\chi_2, \chi_1$ | 0.07,0.055 | 12 | 1.2 | n.a.* | n.a. | n.a. |
| Y23 | $\chi_2$, CMAP | 0.07,0.30 | 12 | 1.2 | 10 | 0.6 | 8 |
| F33 | $\chi_2, \chi_1$ | 0.07,0.07 | 8 | 1.2 | 10 | 0.6 | 12 |
| Y35<sup>c</sup> | $\chi_2, \chi_1$ | 0.07,0.07 | 12 | 1.2 | 12 | 0.6 | 12 |
| F45 | $\chi_2, \chi_1$ | 0.07,0.07 | 12 | 1.2 | 10 | 0.6 | 8 |
| F45 (c36) | $\chi_2, \chi_1$ | 0.07,0.07 | n.a. | n.a. | 12 | 0.6 | 8 |

<sup>a</sup>charmm36/standard TIP3P simulations, <sup>b</sup>charmm36m/charmm TIP3P, <sup>c</sup>non-converged free energy surface. \*n.a.: simulations not performed in this case.

For residue Y23, a contact-map based collective variable was used along with *χ*_2_. This variable is defined using the following function

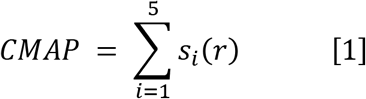

where *s_i_*(*r*) is a switching function for each atom pair considered.

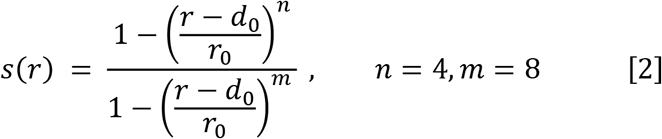

The values of *r*_0_ and *d*_0_ are mentioned in **Table S1 in SI** and are based on equilibrium distances (obtained from unbiased simulation) between *C_α_* atoms of five residue pairs viz. A25-G28, G28-G56, C55-A25, C55-C5, A25-C5. We note that such a contact map based collective variable is heuristic, and it is probably possible to define a better CV. For all WTMetaD simulations, the hill width was chosen as one half of the standard deviation of the CV in a free simulation run. The hills were added at an interval of 1 ps.

In some simulations, repulsive harmonic wall bias was (*V_b_*) imposed to restrict sampling up to specific values of either *χ*_2_ or *χ*_1_ values.

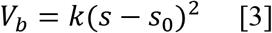

where *k* is the force constant in kJ mol^−1^ rad^−2^, *s*_0_ limiting the value of *χ*_2_ or *χ*_1_ in radians. *UPPER_WALLS* and *LOWER_WALLS* keywords in PLUMED are used to avoid sampling above and below the specific limiting value *s*_0_, respectively. The details of parameters *k* and *s*_0_ are provided in Table S3 in SI.

A time-dependent metadynamics bias *V*(*s, t*) corrected with time-dependent bias offset^35^ *c*(*t*) was used for reweighting^36^. The expression of *c*(*t*) is shown below:

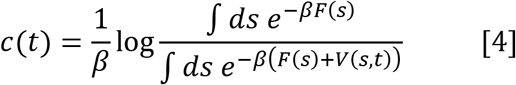

The evolution of *c*(*t*) with simulation time is shown in **figure S1 of SI** to understand the quasi-equilibrated region in WTMetaD simulations. We also tested the variational approach to conformational dynamics^37^ (VAC-MetaD) approach to avoid misleading results due to suboptimal CVs (see Appendix II in SI for further details). The quasi-equilibrated part was utilized in the case of residue F22 and F33 to build CVs using VAC-MetaD^37^.

NMR experiments^38–39^ have observed that the C14 *χ*_1_ motions are 30-fold faster than C38 *χ*_1_ and lead to low populated state *m* and a highly populated state *M*. The disulfide bond orientations arising due to rotations *χ*_1_ of torsion of C14 and C38 residues in BPTI is known as disulfide bond isomerization. The DESRES simulation observed five long-lived BPTI conformational states, with each state having different C14-C38 disulfide bond conformation. The ring-flipping rates were different in each state, suggesting a relation between C14-C38 disulfide bond isomerization and ring-flipping in BPTI. We reweighted the histogram of *χ*_1_(C14)-*χ*_1_(C38) and *χ*_2_(C14)-*χ*_2_(C38) to understand BPTI conformations sampled in converged WTMetaD simulations.

### Estimation of the flipping rate from metadynamics simulation

The reliability of InMetaD depends on two major factors viz. (i) chosen collective variables should discriminate between metastable states, (ii) no bias should be deposited in the transition state (TS) region. The converged free energy surface along biased CVs indicates the sampling along possible slow degrees of freedom, which are the bottleneck for the sampling. The same set of collective variables can be utilized for the InMetaD simulations. The criterion to avoid bias addition at the TS region can be achieved by a slow (infrequent) addition of gaussian bias during metadynamics. In all cases, we have performed InMetaD simulations by biasing the *χ*_2_ torsion first. These simulations allowed us to understand the efficiency of *χ*_2_ alone to capture ring-flipping dynamics. InMetaD approach to obtain kinetic information simulations is still in infancy. Previous studies^41–43^ have demonstrated the efficiency of this approach to study millisecond timescale dynamics in protein-ligand interactions. For each residue, *N* independent simulations were performed to obtain transition times, where *N* = 40 if not otherwise stated. For each i^th^ individual run, the acceleration factor^22^ (*α_i_*) arising due to slow addition of time-dependent bias along CV *s* is given as:

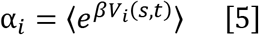

The exponentiated bias is averaged until the time of transition to another basin, i.e., actual simulation time 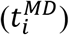, and the transition times (*τ_i_*) of 180° ring flip is obtained as

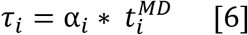

The average transition time (*μ*) from multiple transition times (*τ_i_*) is obtained as:

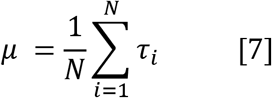

The transition times obtained from InMetaD should follow a Poisson distribution according to the law of rare events. Salvalaglio *et al.^40^* proposed a two-sample Kolmogorov-Smirnov test (KS test) to compare the distribution of transition times (*τ_i_*) to the ideal Poisson distribution. In KS-test, the characteristic ring-flip time (t_flip_) is obtained by fitting the theoretical cumulative distribution function (TCDF) to the empirical cumulative distribution function of InMetaD derived *τ_i_* values.

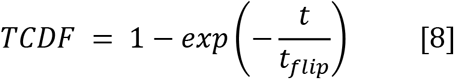

This goodness of fit test helps to understand the deviation of simulated transition times from Poisson characteristics by observing the ratio of average transition time (*μ*) to the median of transition times *τ_i_* (*t_m_*), and the p-value (significance level 0.05). If the *τ_i_* values follow a Poisson distribution; the ratio μ ln 2/t_m_ should be equal to 1. This ratio is sensitive to the deviation from the Poisson distribution and is reported in the present study along with p-values. The inverse of fitted t_flip_ value is considered as the flipping rate (*k_flip_*) as it has been observed to converge faster^40^ than the average (*μ*) in a previous study^40^.

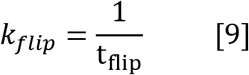

The InMetaD simulations were stopped when the trajectory crosses the barrier and transit to the stable state of interest (i.e., either 180° flipped state or intermediate) by implementing boundary conditions based on biased CV values. The boundary conditions are defined closer to the product using the COMMITOR keyword in PLUMED and not precisely at the speculated transition state CV values. The boundary CV values are monitored at every 10 ps or 50 ps. This protocol assures a complete transition to a flipped state and is not misinterpreted due to recrossing events. In the recrossing events, the trajectory does not continue to the flipped state after crossing a barrier but rather transit back to the starting state minima.

## RESULTS AND DISCUSSION

We characterized aromatic ring flipping in BPTI in two complementary ways. First, we used the well-tempered metadynamics (WTMetaD) simulations to calculate the free-energy surface and determine the most relevant intermediates. Second, we used the InMetaD method to calculate flipping rates and distinguish various mechanisms. An important part of the investigation consisted of choosing suitable collective variables (CVs) to be biased in both types of simulations.

The overall results obtained from the InMetaD investigation of each residue are shown in Table 2, together with the corresponding CVs used. The experimental ring flip rates^3^ (*k_exp_*) at 300 K and rates obtained from 1-ms-long DESRES simulation (*k_des_*) are also shown for comparison. We used the procedure in ref. 3 to calculate *k_exp_* values.

**Table 2.** InMetaD derived ring flip time from exponential fit (t_flip_) at 300 K, ring flip rate (***k***_flip_), KS-test p-value and the ratio of average (μ) to a median of Poisson distribution from InMetaD simulations, experimental ring flip rate (*k_exp_*) at 300 K, and from a 1-ms-long DESRES trajectory (*k_des_*).

| Residue | CVs | $t_{\text{flip}}$ (s) | $k_{\text{flip}}$ ($s^{-1}$) | p-value | $\mu \ln 2/t_m$ | $k_{\text{exp}}$ ($s^{-1}$) | $k_{\text{des}}$ ($s^{-1}$) |
| --- | --- | --- | --- | --- | --- | --- | --- |
| F4 | $\chi_2, \chi_1$ | $2.80 \times 10^{-5}$ | $3.57 \times 10^4$ | 0.32 | 1.79 | $> 10^3$ | $7.0 \times 10^6$ |
| Y10 | $\chi_2, \chi_1$ | $5.80 \times 10^{-5}$ | $1.72 \times 10^4$ | 0.80 | 1.54 | $> 10^3$ | $1.0 \times 10^6$ |
| Y21 | $\chi_2$ | $5.20 \times 10^{-3}$ | $1.92 \times 10^2$ | 0.68 | 1.32 | $< 100^a$ | $1.0 \times 10^3$ |
| F22 | $\chi_2, \chi_1$ | $2.41 \times 10^{-2}$ | $4.15 \times 10^1$ | 0.16 | 3.54 | 670<sup>b</sup> | $7.0 \times 10^4$ |
| F22 (c36)<sup>d</sup> | $\chi_2, \chi_1$ | $5.45 \times 10^{-4}$ | $1.83 \times 10^3$ | 0.69 | 1.24 | 670<sup>b</sup> | $7.0 \times 10^4$ |
| Y23 | $\chi_2$, CMAP | $9.81 \times 10^{-1}$ | 1.02 | 0.09 | 1.80 | 140<sup>c</sup> | $1.0 \times 10^3$ |
| F33 | $\chi_2, \chi_1$ | $3.50 \times 10^{-5}$ | $2.86 \times 10^4$ | 0.03 | 2.47 | $> 10^3$ | $6.0 \times 10^6$ |
| Y35 | $\chi_2, \chi_1$ | 3.49 | $2.87 \times 10^{-1}$ | 0.58 | 3.22 | 4<sup>c</sup> | $1.0 \times 10^4$ |
| F45 | $\chi_2, \chi_1$ | 3.22 | $3.11 \times 10^{-1}$ | 0.95 | 1.44 | 220<sup>c</sup> | 0 |
| F45 (c36)<sup>e</sup> | $\chi_2, \chi_1$ | $1.78 \times 10^{-1}$ | 5.61 | 0.97 | 1.40 | 220<sup>c</sup> | 0 |

<sup>a</sup>308 K, <sup>b</sup>278 K, <sup>c</sup>values estimated using the Arrhenius equation and with data from Ref. <sup>3</sup>, results from <sup>d</sup>16 simulations and <sup>e</sup>12 simulations.

### Free Energy and Flipping Mechanism

The results for all eight residues are mentioned in this section. For all free energy profiles, the position of the starting configuration is denoted with state A and intermediates with In (n=1,2). The corresponding symmetry-equivalent states are indicated with prime notation (A’ and I_n_’, n=1,2). Representative structures of A and I_n_ states are shown for each residue. The convergence of metadynamics simulations and diffusion along CVs for each simulation are shown in supporting information (**figures S2 to S10**). The distributions of InMetaD trajectory endpoints immediately after flipping (started from state A) in each case are shown in figures **S11 and S12 in SI**. Following the NMR-motivated “fast” and “slow” classification of aromatic ring flips, the fast-flipping residues (F4, F33, and Y10) are described first, and the remaining slow flipping residues are described later. However, note that the fast-flipping rings in the BPTI exhibit t_flip_ values on the microsecond time scale, making it challenging to obtain statistically significant results. Residues F4 and Y10 (among other residues) could flip via multiple pathways; the t_flip_ values and KS-test derived parameters for each mechanism are mentioned in **Table 3.**

**Table 3.** Mechanism Dependent Results of the Infrequent Metadynamics Simulations of Residue F4 and Y10.

| Residue | mechanism | $N_{\text{traj}}^{\text{a}}$ | $t_{\text{flip}} (s)$ | $k_{\text{flip}} (s^{-1})$ | p-value | $\mu \ln 2/\text{tm}$ |
| --- | --- | --- | --- | --- | --- | --- |
| F4 | All | 40 | $2.80 \times 10^{-5}$ | $3.57 \times 10^4$ | 0.32 | 1.79 |
| | $A \rightleftharpoons I1 \rightleftharpoons I1' \rightarrow A'$ | 22 | $3.02 \times 10^{-5}$ | $3.31 \times 10^4$ | 0.82 | 1.48 |
| | $A \rightleftharpoons I1 \rightarrow I2'$ | 6 | $2.21 \times 10^{-5}$ | $4.52 \times 10^4$ | 0.71 | 0.80 |
| | $A \rightarrow A'$ | 12 | $2.81 \times 10^{-5}$ | $3.56 \times 10^4$ | 0.13 | 2.78 |
| Y10 | All | 40 | $5.80 \times 10^{-5}$ | $1.72 \times 10^4$ | 0.80 | 1.54 |
| | $A \rightleftharpoons I1 \rightarrow I1'$ | 36 | $5.79 \times 10^{-5}$ | $1.73 \times 10^4$ | 0.89 | 1.54 |
| | $A \rightarrow A'$ | 4 | $3.42 \times 10^{-5}$ | $2.92 \times 10^4$ | 0.39 | 1.48 |

<sup>a</sup> $N_{\text{traj}}$ = number of trajectories following a particular mechanism.

### Fast flipping residues

#### Residue F4

Residue F4 is located on the solvent-exposed protein exterior and has frequent interactions with residue R42. The free-energy surface is shown in Figure 2 together with representative structures. State A’ and state I2’ are considered “flipped” states, signaling that transition has occurred, thus providing *τ_i_* (*i* = 1 *to* 40) values. The results are shown in **figure S13** and the bootstrap analysis of all 40 *τ_i_* values and the p-value obtained from the KS test are shown **in figure S14 in SI.** The effective t_flip_ considering all 40 simulations was *2.80 ×* 10^−5^ s.

**Figure 2.**
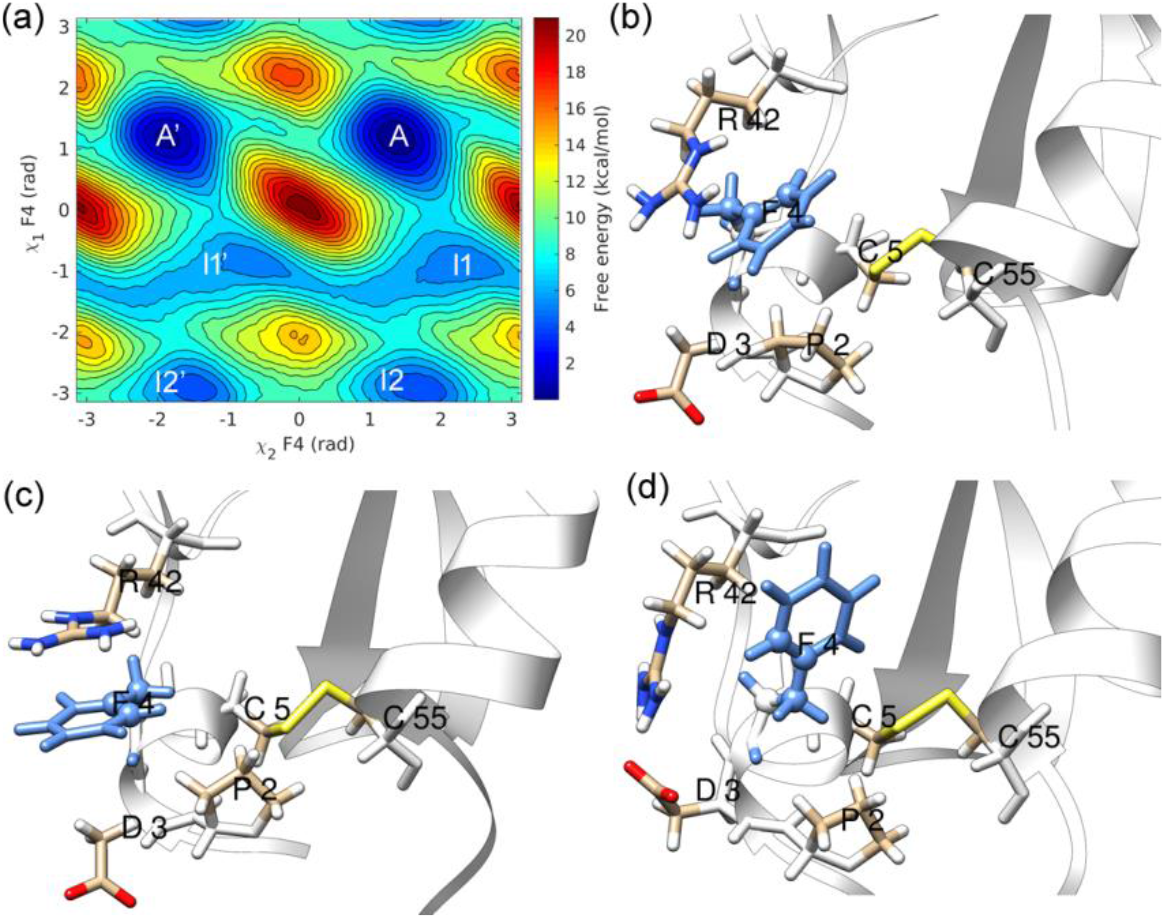
(a) Residue F4 (blue) ring-flipping free energy along *χ*_2_ and *χ*_1_ reaction coordinates and representative structures from (b) minimum A, (c) minimum I1, and (d) minimum I2.

State I1 is involved in the ring-flipping of residue F4. The cation-π interactions with positively charged residue R42 stabilizes the conformations in minimum I1. Residue F4 is located close to the disulfide bond [5-55] formed by C5 and C55. The disulfide bond [5-55] torsion was not affected during states A to I1 transition because the aromatic side-chain of F4 moved away from residue C5. State I2 has a trans orientation of the bonds N-CA (backbone) and CB-CG (side-chain), thus exposing R42 towards solvent. Salt bridge interactions were observed between R42 and D3 in state I2.

Three different mechanisms were observed during the InMetaD simulations, as described in Table 3. A to A’ via intermediates is the prominent mechanism observed during F4 ring-flips. In this mechanism, ring-flips occur via coupled motions of the *χ*_2_ and *χ*_1_ torsions, starting from state A, ring transit to state I1 multiple times, which acts as a steady-state before reaching state I1’ and eventually proceed to A’. A direct ring flip from state A to A’ was also observed, but these trajectories exhibited a few A to I1 reversible transitions before the direct transition. Finally, six trajectories flipped to state I2’ while started from state A. Reversible transitions between states A and I1 were observed due to motions along *χ*_1_. These trajectories represent an off-pathway mechanism where trajectories before proceeding to state I1’ transition to state I2’, thus achieving flipped orientation. Therefore, multiple flipping pathways are possible with almost the same t_flip_ in case of residue F4.

#### Residue Y10

We have observed two distinct mechanisms for flipping: a *χ*_2_ − *χ*_1_ coupled mechanism and a direct ring flip. The prominent mechanism of residue Y10 involves motions along the *χ*_1_ torsion leading to stable state I1 (see figure 3(a)). In that state, residue Y10 is less restricted by interactions with surrounding atoms (see fig. 3(c)) as the motions along *χ*_1_ brings Y10 aromatic side-chain out of the core region surrounded by T11, K41, and P13. We have observed transient interactions with K41 but no hydrogen bonding interactions. The aromatic ring-flipping in Y10 illustrates an alternate mechanism to the “breathing cavity” mechanism proposed by earlier studies. This alternate mechanism involves moving the aromatic side-chain to the less crowded region and undergoing the flipping process. This mechanism is different from the “cavity model,” where transient cavity formation allows ring-flipping.

**Figure 3.**
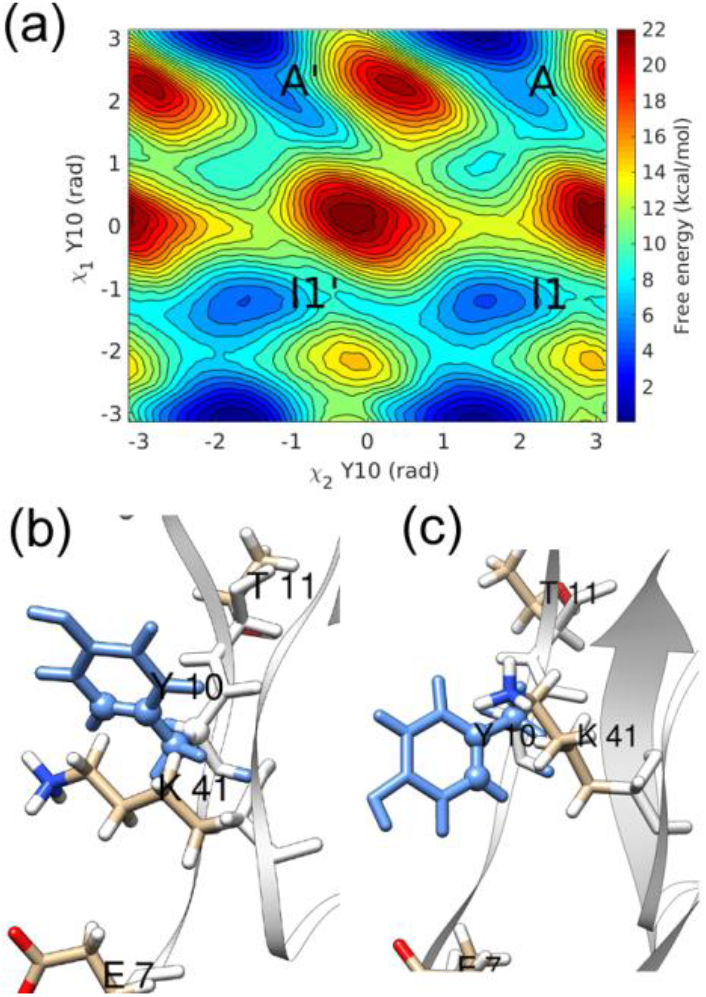
*χ*_2_ − *χ*_1_ free energy surface, and representative structures in (b) minimum A and (c) minimum I1 of the free energy landscape of residue Y10 (blue).

Most InMetaD trajectories (36 out of 40 simulations) followed the *χ*_2_ − *χ*_1_ coupled mechanism involving state I1. A *t_flip_* of 5.80 × 10^−5^ *s* was observed by considering all 40 InMetaD simulations. The KS-test results and associated errors are reported in **figures S15 and S16 in SI.** Four trajectories have transitioned to flipped state *A’*. However, there were a few (one or two) A-I1 jumps that occurred before proceeding directly to state A’. Thus, state I1 is an important intermediate that could be an active or off-pathway state depending on the transition path.

#### Residue F33

The residue F33 is surrounded by hydrophobic residues P9, F22, and Y35. The significant variation in values of *χ*_1_ torsion during WTMetaD simulations leads to states I1 and I2, where the aromatic side-chain of F33 is out of the protein interior (see figure 4). The flipping of F33 only required small fluctuations relative to the backbone. Thus, even though states I1 and I2 were observed in the free energy landscape, the direct ring-flipping pathway did not involve these states. All trajectories visited an off-pathway state I1 during the InMetaD simulations. The F33 flipping affects *χ*_1_ fluctuations of residue F22 (**see figure S17 in SI**). All InMetaD simulations showed flipping mainly by *χ*_2_ rotation with positive values of *χ*_1_ (**figure S11 in SI**). A *t_flip_* of 3.50 × 10^−5^ s was obtained through the KS test and p-value of 0.03. To understand the small p-value, indicating significant deviation from the Poisson distribution, we analyzed the InMetaD trajectories and found deposition of bias in the barrier region along *χ*_1_ (*χ*_1_ ≈ 2.4 rad). The example of bias deposition in randomly chosen trajectories is shown in **figure S18** in the supporting information. Residue F33 demonstrated nonproductive off-pathway transitions between A and I1 states, whereas the main mechanism is direct flipping. To investigate whether these non-productive pathways affected the results, we performed another set of 12 InMetaD simulations, in which a repulsive wall bias restricted the sampling of *χ*_1_ values greater than 2.4 rad (see method section). The *t_flip_* value from 12 simulations with wall bias is close to previous *χ*_2_ − *χ*_1_ InMetaD simulations without wall bias (**see Table S2 and figures S19-S20 in SI**).

**Figure 4.**
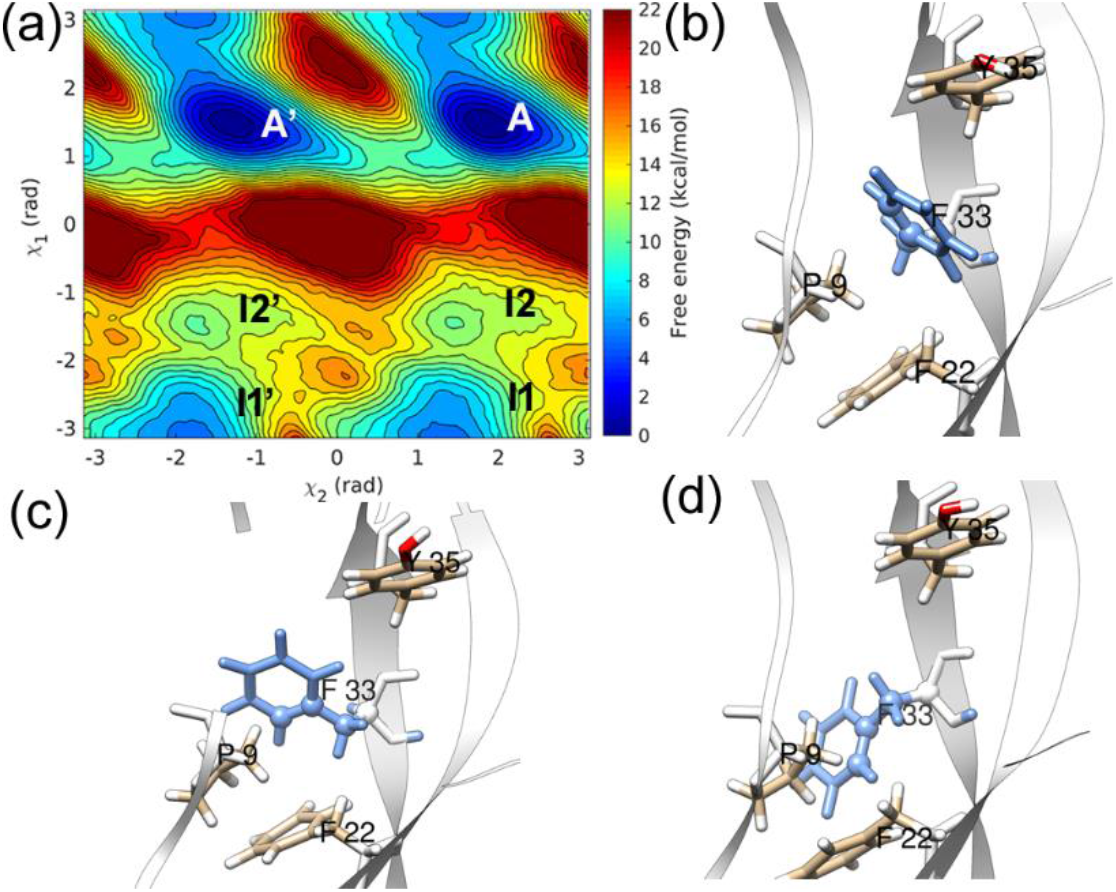
(a) *χ*_2_ − *χ*_1_ free energy surface, and representative structures in (b) minimum A, (c) minimum I1, and (d) minimum I2 of the free energy landscape of residue F33 (blue).

### Slow flipping residues

#### Residue Y21 and F45

As shown in Figure 5, residues Y21 and F45 both lack intermediates in the free energy surface and flip via a simple direct ring flip mechanism. Residue Y21 displayed convergence when only the *χ*_2_ torsion was used as a CV and the resulting *t_flip_* = 5.20 × 10^−3^ s would classify it as a slow-flipping residue. Residue Y21 is solvent-exposed, and thus simple motions along *χ*_2_ torsion is sufficient as no steric hindrance with the surrounding protein matrix occurs. Instead, most of the ring flips occur in a narrow range of *χ*_1_ values. The reweighted distribution of *χ*_2_ − *χ*_1_ from the converged metadynamics simulation indicates that no significant motions along *χ*_1_ is needed, and Y21 can flip transiently. This residue lies on the rigid β-strand (I18-N24), which forms an anti-parallel β-sheet via strong interactions with another β-strand (L29-Y35). Interestingly residue F33 also lies on a β-strand but is a fast-flipping residue.

**Figure 5.**
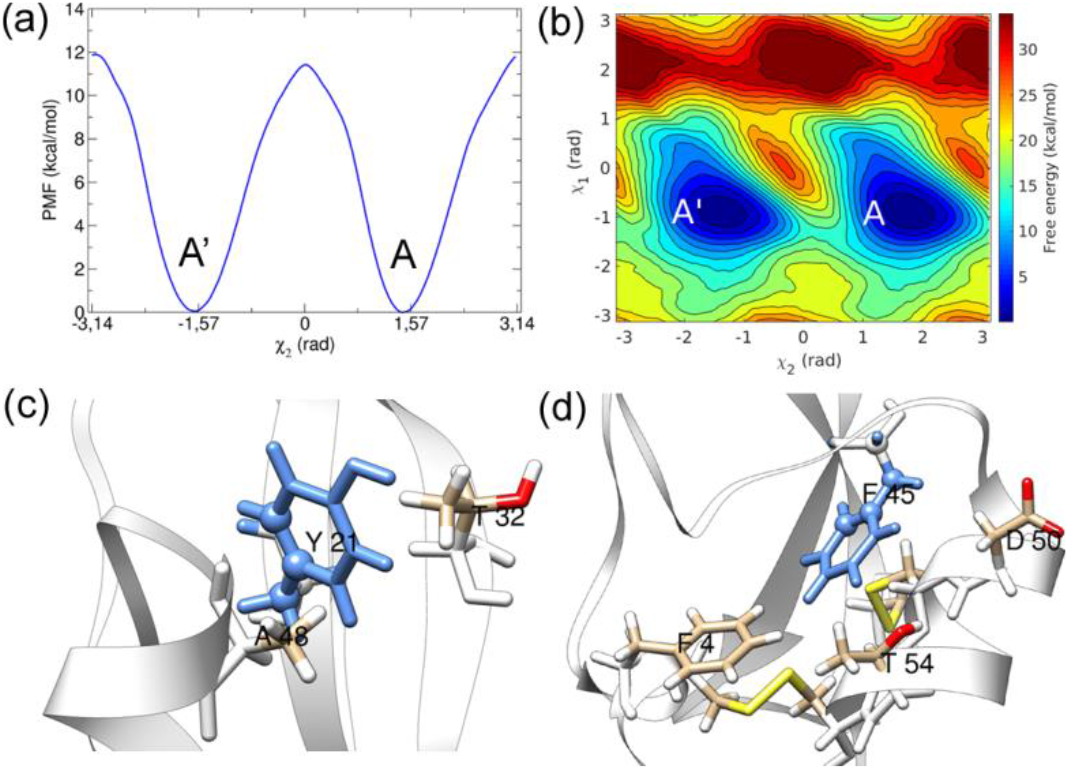
Free energy of (a) residue Y21 (blue) along *χ*_2_, (b) residue F45 (blue) along *χ*_2_ − *χ*_1_, (c) representative structure from minimum A of residue Y21, and (d) from minimum A of residue F45.

A converged free energy surface was observed along *χ*_2_ − *χ*_1_ collective variables for residue F45, and thus, 40 InMetaD simulations were performed along the *χ*_2_ − *χ*_1_ variables. The distribution of *τ_i_* (i=1 to 40) passes KS-test and fitted *t_flip_* of 3.21 s was obtained, indicating F45 as a slow flipping residue. There is direct flipping in F45; however, *χ*_1_ is needed for librational motions (see Fig. 5(d)), which facilitate flipping along *χ*_2_ (*InMetaD simulations based on *χ*_2_ did not pass the KS-test as discussed below*). The cumulative distribution function of *τ_i_* and associated errors are shown in **figures S21 and S22 in SI.**

#### Residue F22

Figure 6 shows the *χ*_2_ − *χ*_1_ free energy surface for F22 along with representative structures. All the InMetaD simulations followed a direct A → A’ transition pathway as a high barrier exist along *χ*_1_ to visit intermediate states. t_flip_ of 2.41 × 10^−2^ s was observed for a direct transition. Direct flip mechanism suggests residue F22 needs minimal fluctuations relative to the backbone for ring-flipping and it might not be necessary to span the whole *χ*_1_ torsion range, as observed in WTMetaD simulations. State I1 occurs due to changes *χ*_1_ value and transient increase of the P9-F33 distance. In state I2, there is no stacking of F22 over F33, but these residues align approximately parallel to each other. During a long 1.6 μs WTMetaD simulation, we have observed a single ring flip event of residue F33 (**figure S23 in SI**). The KS-test results for residue F22 are reported in Figures **S24 and S25 in SI.**

**Figure 6.**
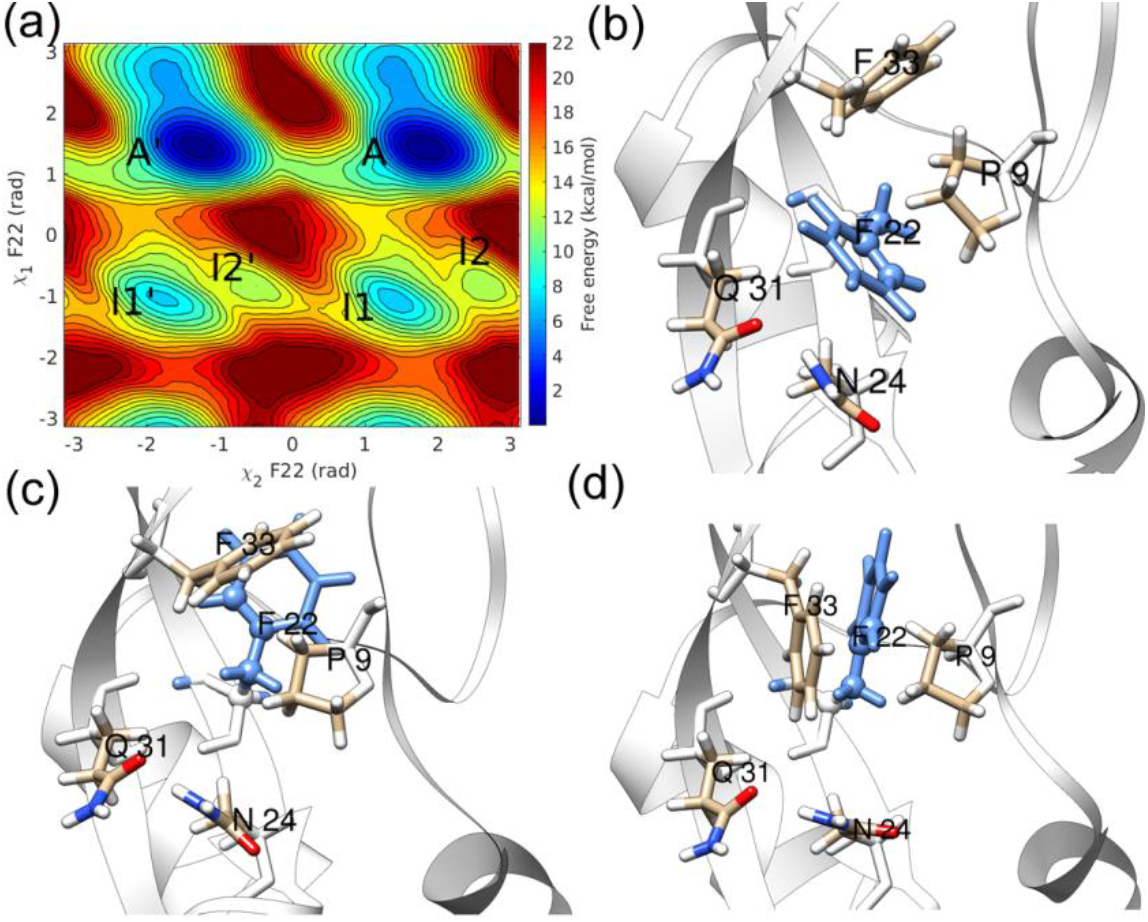
(a) Free energy of residue F22 (blue) along *χ*_2_ − *χ*_1_ and representative structure from (b) minimum A, (c) minimum I1, and (d) minimum I2.

**Figure 7.**
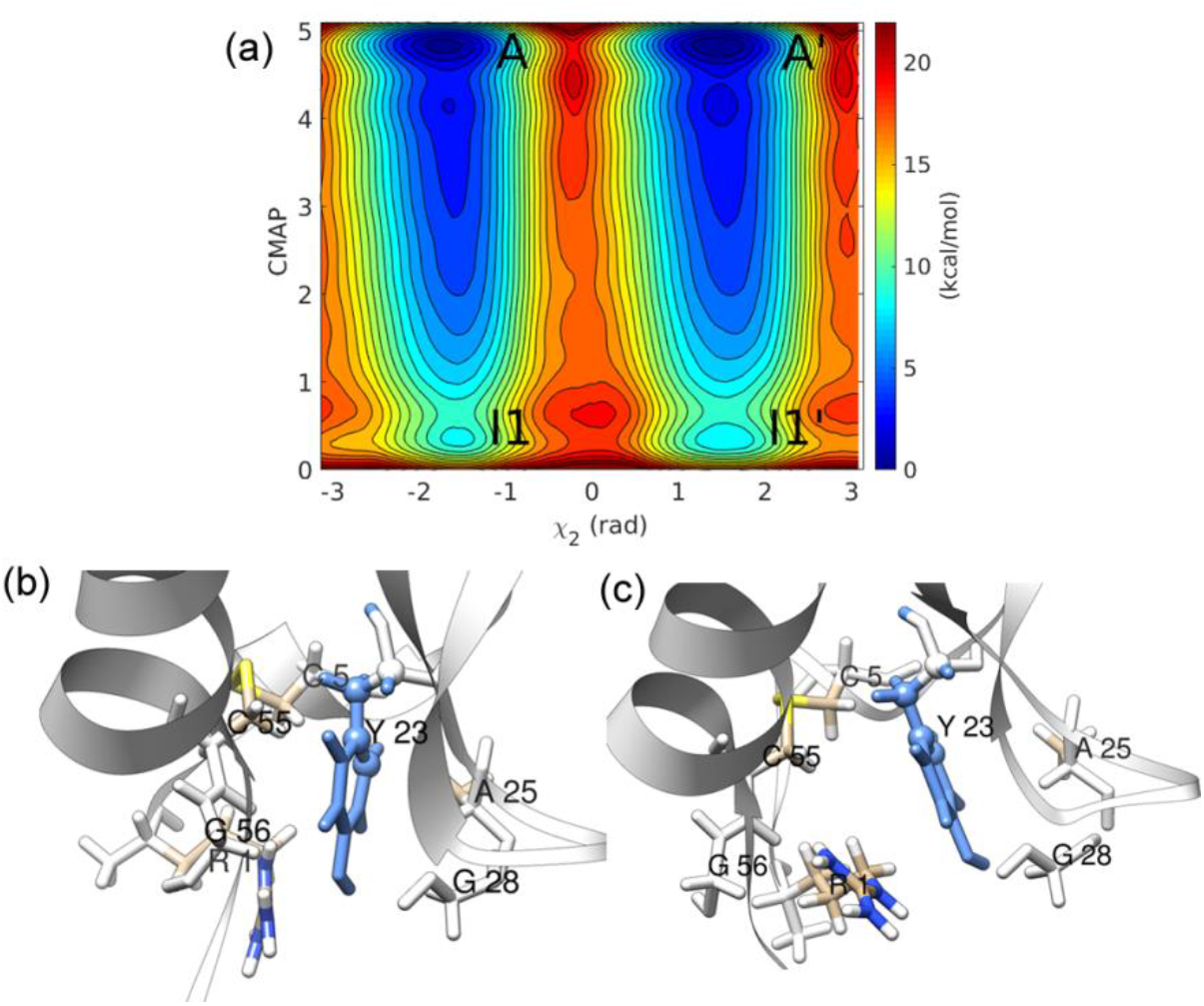
Free energy of (a) residue Y23 along *χ*_2_ − *CMAP*, and representative structure from (b) minimum A, and (c) minimum I1.

Five InMetaD simulations in each case of (a) A → I1 pathway, (b) I1 → A pathway, and (c) I1 → I1’ pathway were performed. In these simulations, the visits to other stable states were avoided using repulsive wall bias^41^ using *k* and *s*_0_ mentioned in **Table S3 in SI.** The results (**Table S4 in SI**) and details of these simulations are reported in **Appendix I in SI**. State-to-state transition rate matrix is reported in **Table S5 in SI**. The dominant eigenvalue 83 s^−1^ was estimated by solving this transition matrix.

#### Residue Y23

Residue Y23 is one of the slowest flipping residues. The dynamics of ring Y23 was studied using the *χ*_2_ torsion and a contact map of C_α_ atoms surrounding the ring (**figure S26 in SI**). The convergence in WTMetaD simulations and KS test validation assured the choice of CMAP as a CV complementary to *χ*_2_. The transition is possible at CMAP values between 1 and 4, indicating that a small change in the equilibrium distances of C_α_ atoms is sufficient to provide the necessary volume for ring-flipping. Thus, a transient excited state could exist during ring-flipping, in which the surrounding atoms are not significantly displaced. We have performed 60 InMetaD simulations to obtain reliable statistics, and observed a *t_flip_* of 0.98 s (see **Figures S27 and S28 in SI**).

Residue Y23 is located near two disulfide bridges defined by residues 5-55 and 30-51. The aromatic side-chain of Y23 infrequently interacts with C_α_, S_γ_, N atom of residue C30 and backbone C, O atoms of residue C5. However, no significant changes in the values of *χ*_2_, *χ*_1_ and *χ*_3_ torsions of the 5-55 and 30-51 disulfide bridges were observed (**figure S29 in SI**). Transient hydrogen bonds between residue Y23 and residues R1, G56, D3 were observed (**figure S30 in SI**). The fluctuations in the flexible residue R1 also transiently contributed to the cavity expansion due to interactions with residue G56. We have also performed *χ*_2_ − *χ*_1_ biased WTMetaD simulations; however, we observed a lack of convergence (**figures S31 and S32 in SI**). Visual observation of the WTMetaD simulation indicated a crowding effect of neighboring C_α_ atoms. Thus, a contact map was used as a complementary CV to obtain a converged free energy landscape. Both *χ*_2_ − *χ*_1_ (t_flip_ = 0.61 s) and *χ*_2_ − *CMAP* (t_flip_ = 0.98 s) InMetaD simulations provided similar t_flip_ values.

Karplus and Gelin^42^ theoretically studied aromatic ring-flipping for Y10, Y21, Y23, and Y35 residues of BPTI by varying the *χ*_1_ and *χ*_2_ torsions. The study concluded that a few strong nonbonded interactions with surrounding atoms lead to a specific ring-flipping rate constant (k_flip_) for each residue. In the DESRES simulation, the ring-flipping rate was not correlated with the density of heavy atoms within 4 Å of aromatic ring atoms at δ and ε positions^19^. This lack of correlation suggests no significant displacement of atoms surrounding the aromatic residue, contrary to the well-accepted cavity model.^12^ We speculate that small displacements of a few surrounding atoms are sufficient to allow ring-flipping. The results for residue Y23 in this work support this hypothesis.

#### Residue Y35

Y35 is a slow flipping residue in BPTI and is one of the most studied aromatic side-chains by previous^10, 43–46^ theoretical studies. The lack of converged free energy profiles for Y35 with *χ*_2_ as well as *χ*_2_ − *χ*_1_ biased metadynamics simulations indicates a missing orthogonal degree of freedom and probably a complex ring-flipping mechanism. Residue Y35 is located inside the protein core, close to the C14-C38 disulfide bond (figure 8), and near the initial part of the loop region formed by residues G36-A40. WTMetaD simulations along *χ*_2_ − *χ*_1_ (1.6 μs) and *χ*_2_ (0.7 μs) for residue Y35 lacked convergence (**figures S7 and S8 in SI**). Thus, we speculate that more than two descriptors might be needed to understand the flipping of the Y35 aromatic side-chain.

**Figure 8.**
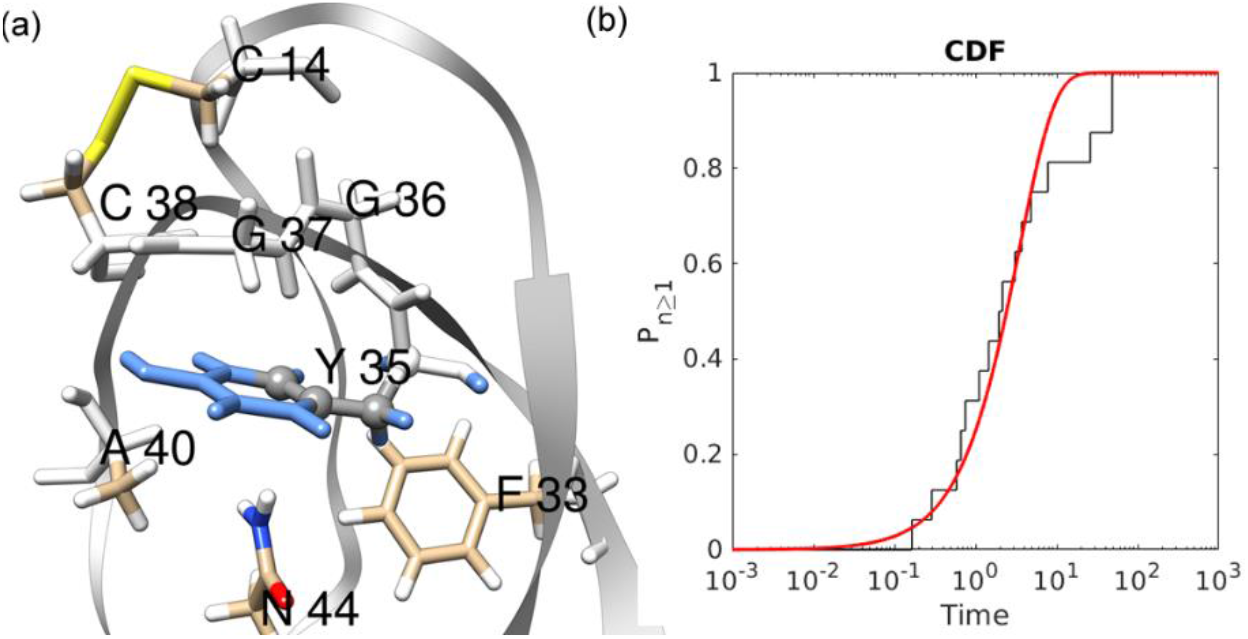
(a) Representative structure of residue Y35 in the native state, (b) Poisson fit to CDF of transition timescales obtained from *χ*_2_ − *χ*_1_ InMetaD simulations (p-value = 0.58 and μ ln 2/tm = 3.21).

The reweighted distribution of side-chain torsion *χ*_1_ of C14 and C38 residues indicate that this simulation has sampled more than state M (**figure S33 in SI**). Thus, the loop region’s conformations due to C14-C38 disulfide bond motions could affect the flipping rate.^19^ We performed 16 InMetaD simulations, assuming *χ*_2_ − *χ*_1_ could provide rates in major state M1 and *t_flip_* of 3.4 s was obtained, indicating Y35 as a slow flipping residue.

The disulfide bond isomerization could likely be associated with non-convergence of the *χ*_2_ − *χ*_1_ metadynamics simulations. Even in a 1-ms-long trajectory, multiple flipping rates of residue Y35 were reported based on C14-C38 disulfide bond isomerization^19^. Thus, we have also performed WTMetaD simulations with *χ*_2_ and *χ*_1_ of Y35, together with the *χ*_3_ dihedral of the C14-C38 disulfide bond. This simulation demonstrated a free energy barrier of 6 kcal/mol for Y35 ring-flipping, as the Y35 aromatic side-chain is irreversibly expelled out of the protein core and was freely rotating in the solvent-exposed state (**figure S34 in SI**).

### Recrossing events F22 and F33 ring-flipping

*χ*_2_ − *χ*_1_. biased InMetaD simulations of residues F22 and F33 demonstrated recrossing events. In figure 9, we have presented one such trajectory of residue F33 to understand the structural details of recrossing. In this trajectory, the first attempt of ring-flipping occurred around 22538 ps (figure 9(b)), when the *χ_2_* value reached −2.06 rad while maintaining positive *χ*_1_ values (state R).

**Figure 9.**
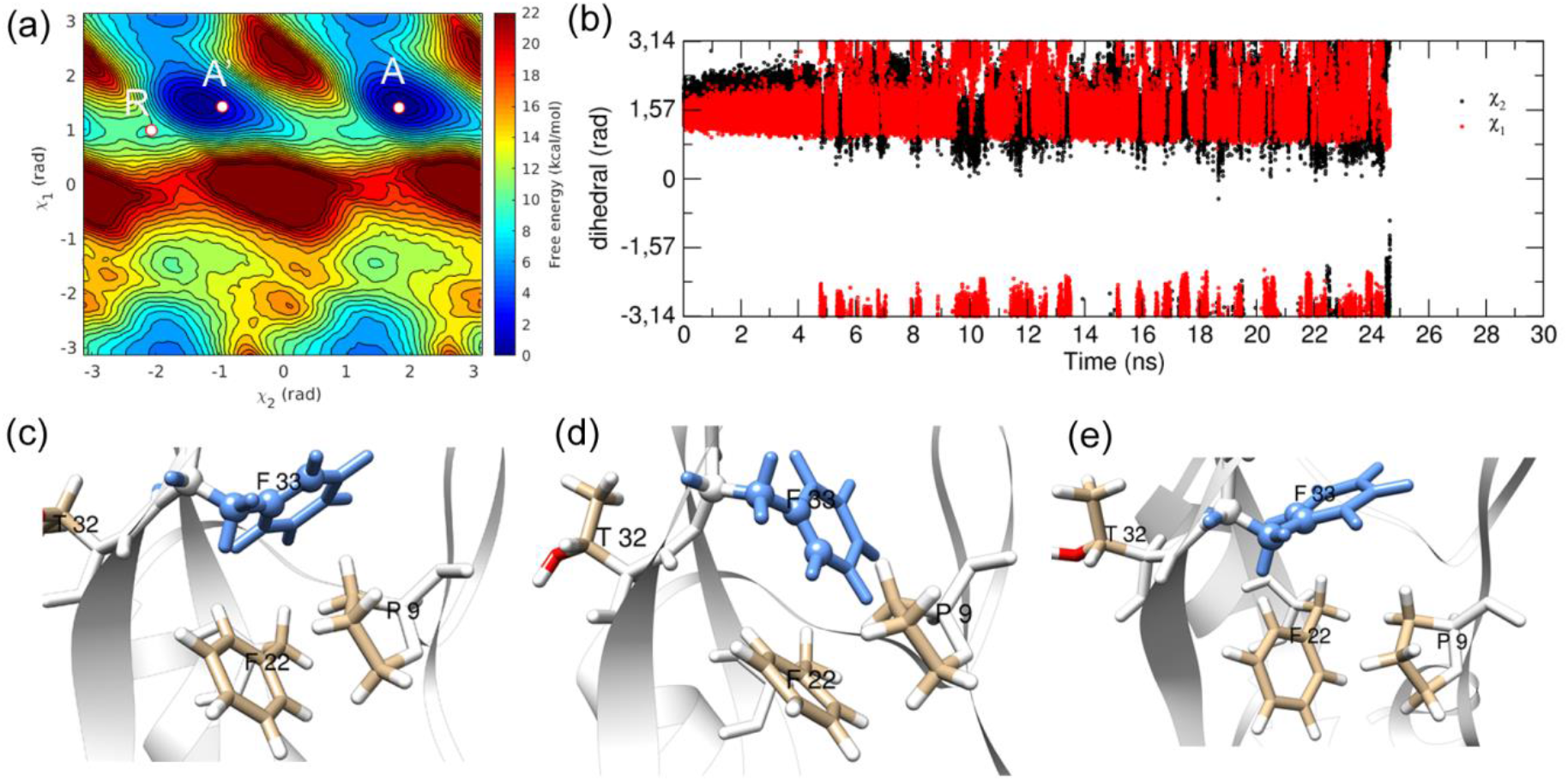
Representative InMetaD trajectory showing recrossing event in F33 ring-flipping. (a) the starting state *A*, flipped state *A’*, and point of recrossing *R* on the free energy surface of *χ*_2_ − *χ*_1_, (b) time evolution of *χ*_2_ and *χ*_1_ torsion angles, (c) orientation of F33 in the native state(state *A, χ*_2_ = 1.83 rad, *χ*_1_ = 1.41 rad), (d) structure at 22538 ps (state *R, χ*_2_ = −2.06 rad, *χ*_1_ =0.99 rad), (e) flipped state at 24643 ps (state *A’*, *χ*_2_ =-0.95 rad, *χ*_1_ = 1.42 rad).

**Figure 10.**
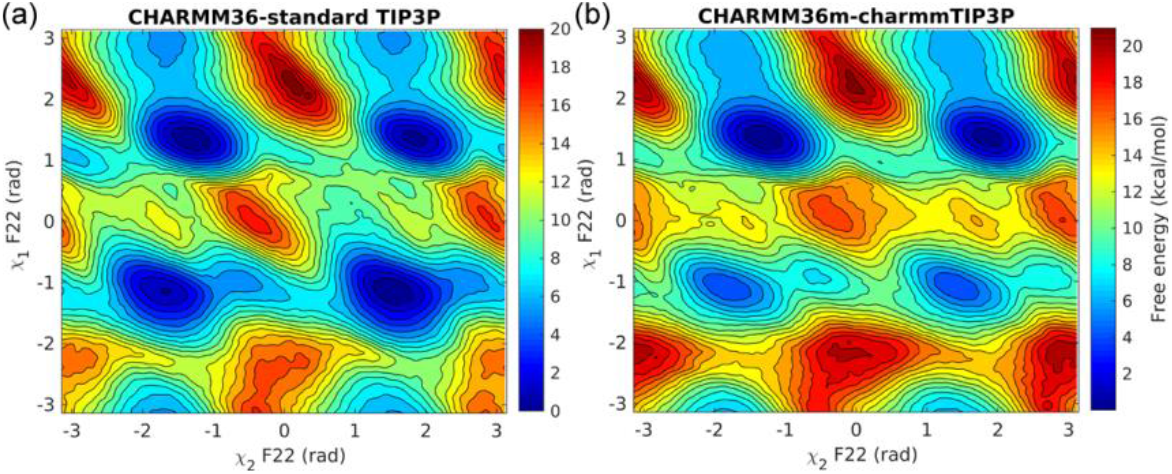
(a) The free energy landscape of residue F22 along *χ*_2_ − *χ*_1_ collective variables with (a) CHARMM36 and TIP3P water model and (b) CHARMM36m and modified TIP3P water model.

*χ*_2_ − *χ*_1_. biased InMetaD simulations of residues F22 and F33 demonstrated recrossing events. In figure 9, we have presented one such trajectory of residue F33 to understand the structural details of recrossing. In this trajectory, the first attempt of ring-flipping occurred around 22538 ps (figure 9(b)), when the *χ*_2_ value reached −2.06 rad while maintaining positive *χ*_1_ values (state R). The ring rotates back to positive *χ*_2_ values after this state and keeps sampling in the initial region with a few *A-I*1 transitions. Transition state theory^47–48^ (TST) assumes that the trajectories cross the barrier region (which divides the reactant and product states) and continue to the product state. TST-based transition rate (k_TST_) can be calculated from the free energy of activation (Δ*G*^‡^) using the Eyring equation [10].

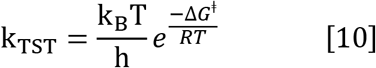

where k_B_ is Boltzmann’s constant, T is the temperature, h is Planck’s constant, and *R* is the gas constant. The TST-based rate due to recrossing events is corrected using the transmission coefficient (*κ*):

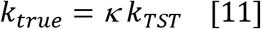

The origin^48–49^ of recrossing events could be associated with the efficiency of collective variables to describe the transition state or with strong coupling between system coordinates undergoing transition and the surrounding environment. However, it is complicated to define the precise TS. Thus, according to variational TST^48–49^, an optimized reaction coordinate that minimizes recrossing events should be defined to understand dynamics. For residue F22, a barrier of 12 kcal/mol exists along the direct flipping pathway, thus estimating the ring-flipping rate, *k_TST_* = 1.13 × 10^4^ s^−1^, according to equation [10]. If collective variables *χ*_2_ and *χ*_1_ are speculated to capture the dynamics of F22 ring-flipping correctly, then the true rate of ring-flip (*k_true_*) is the same as the one obtained from InMetaD simulations (k_InMetaD_) i.e. *k_true_* = k_InMetaD_. The substitution of *k_TST_* and k_InMetaD_ values in equation [11] then suggests that *κ* = 3.67 × 10^−3^.

Based on the variational TST assumption, this unusually low transmission coefficient *κ* value could be attributed to the choice of CVs, *χ*_2_ and *χ*_1_, used for InMetaD simulations. To verify this assumption, we performed 12 InMetaD simulations in each case of residue F22 and F33 using reaction coordinates optimized using a variational approach (VAC-MetaD).^37^ This approach allows a linear combination of multiple descriptors of the process and enables a better description of transition states and stable states. We have considered the distance between residues P9 and F33, *χ*_1_ and *χ*_2_ torsions of residues F22 and F33 to build VAC-optimized CVs (VAC-CVs). The computational details and results of these simulations are reported in the SI (**Appendix II in SI**).

With VAC-CVs, *t_flip_* values 0.09 s (p-value = 0.49) and 8.64 × 10^−5^ s (p-value=0.70) were obtained for residues F22 and F33, respectively (**figure S47 in SI**). Thus, even with optimized CVs, residue F22 gave a low κ value. Furthermore, we have also observed recrossing events in a few F22 and F33 VAC-CV based simulations. Thus, further analysis is needed to understand the origin of such low transmission coefficient and the corresponding frictional effects. A recrossing event was also observed for c36 based InMetaD simulations of F22.

#### Limitations of 1D *χ*_2_ InMetaD simulations

We performed InMetaD simulations biasing *χ*_2_ as a standalone CV to understand its limitations and accuracy to estimate t_flip_ values. The details of metadynamics parameters and the outcome of *χ*_2_ biased InMetaD runs are reported in **Table 4**. KS test results for all residues are reported in **figures S39 and S40 in SI.** *χ*_2_ biased InMetaD simulations were not performed for residue Y35, as WTMetaD simulations biasing *χ*_2_ of Y35 lacked frequent ring-flip events (**figure S8 in SI**).

**Table 4.** *χ*_2_ biased InMetaD derived ring flip rate (***k_χ_2__***,) at 300 K, number of simulations (N), KS-test derived the ratio of average (μ) to the median of Poisson distribution and p-value, InMetaD simulation parameters hill height (w, kJ.mol^−1^), bias factor (*γ*), frequency of hill addition (*v*, ps).

| Residue | $k_{\chi_2}$ ($s^{-1}$) | N | $(\mu \ln 2)/t_m$ | p-value | $\gamma$ | w | v |
| --- | --- | --- | --- | --- | --- | --- | --- |
| F4 | $3.44 \times 10^4$ | 12 | 2.98 | 0.86 | 15 | 1.2 | 12 |
| Y10 | $5.88 \times 10^2$ | 40 | 1.79 | 0.21 | 10 | 0.6 | 18 |
| Y21 | $1.92 \times 10^2$ | 40 | 1.32 | 0.68 | 10 | 0.6 | 18 |
| F22 | 3.65 | 20 | 2.94 | 0.45 | 8 | 1.2 | 12 |
| Y23 | $4.0 \times 10^{-2}$ | 12 | 3.71 | 0.02 | 12 | 1.2 | 12 |
| F33 | $3.4 \times 10^4$ | 15 | 3.46 | 0.90 | 12 | 0.6 | 10 |
| F45 | $1.2 \times 10^{-1}$ | 40 | 5.39 | 0.00 | 10 | 0.6 | 18 |

*χ*_2_ biased WTMetaD simulations for all residues, except Y21, lacked convergence. An example is shown in **figure S41 in SI** for residue Y10. The slow diffusion of *χ*_2_ and *χ*_1_ of residue Y10 in *χ*_2_ biased WTMetaD simulation suggests that biasing of both CVs is necessary. InMetaD simulations along *χ*_2_ for residue Y10 yielded t_flip_ values two orders of magnitude slower than *χ*_2_ − *χ*_1_ InMetaD. We observed that most of these trajectories had a direct transition from state *A* to *A’* with no significant fluctuations along the *χ*_2_ torsion. This result shows that biased CVs direct the reactive flux along pathways specific to them and could provide different timescales. *χ*_2_ biased InMetaD simulations demonstrate that t_flip_ could be overestimated if diffusion along other relevant degrees of freedom is neglected in InMetaD. It is noteworthy that both *χ*_2_ and *χ*_2_ − *χ*_1_ biased InMetaD simulations of residue Y10 passed the KS-test and has low (*μ*ln2)/*t_m_* ratio. In general, missing descriptors of the process could affect the metadynamics derived t_flip_ values even if the two-sample KS test is passed. Theoretically, multiple CVs are needed to capture all metastable states on the free energy landscape. Thus, recent machine learning-based methods to define a combination of multiple collective variables could be specifically useful to obtain reliable rates with InMetaD.

For residue F22, the InMetaD simulations with the *χ*_2_ torsion passed the KS test but prediected tflip an order of magnitude slower than the *χ*_2_ − *χ*_1_ InMetaD simulations. Interestingly, both types of simulations *χ*_2_ and *χ*_2_ − *χ*_1_ InMetaD) gave much higher t_flip_values than observed in the DESRES trajectory. For residue F45, distribution of *τ_i_* values from the *χ*_2_ InMetaD simulations (**figure S40 in SI**) indicated deviation (p-value of 0.00) from ideal Poisson distribution (although the t_flip_ value obtained from these simulations is comparable with the value obtained from *χ*_2_ − *χ*_1_ InMetaD simulations). This exemplifies that apart from collective variables that discriminate stable states, the CVs which promote the diffusion of such principal CVs are needed for reliable estimation of dynamics from metadynamics.

### CHARMM simulations

We have performed metadynamics simulations using two CHARMM36 force field variants, c36, and c36m, as described in the section, for residue F22 with *χ*_2_ and *χ*_1_ collective variables. The free energy barriers along *χ*_1_ are 3 kcal/mol lower compared to the AMBER ff14SB force field. The convergence of these simulations is reported in SI (**figures S42 and S43**).

The ring flip timescales were estimated for residues F22 (16 simulations) and F45 (12 simulations) modeled with the c36 force field. For residues F22 and F45, t_flip_ values of 5.45 × 10^−4^ s (k_fiip_ = 1.83 × 10^3^ s^−1^, p-value=0.69) and 1.78 × 10^−1^ s (k_flip_ = 5.61 s^−1^, p-value = 0.97) were observed, respectively, predicting slow flipping residues(**figure S44 in SI**), but faster by 1-2 orders of magnitude than AMBER ff14SB. The barrier of 9.0 kcal/mol for F22 suggests k_TST_ = 1.74 × 10^6^ s^−1^, thus giving (again) a low *κ* value of 1.04 × 10^−3^ for the c36 based F22 InMetaD simulations.

## CONCLUSIONS

In the present work, we describe a metadynamics based approach to calculate the ring flipping rate and mechanism for eight individual aromatic rings in BPTI. We utilized the InMetaD method to obtain first passage times (*τ_i_*) to estimate *t_flip_*. This approach successfully distinguished between experimentally categorized “slow” and “fast” residues with an appropriate choice of CVs. Our InMetaD simulations categorized residues F4, F33, Y10 as fast flipping and residues Y21, F22, Y23, F45, and Y35 as slow flipping residues. Resides Y35, Y23, and F45 exhibited timescales up to seconds, significantly slower than millisecond brute-force simulation. The rank order observed for ring flips from slow to fast is as follows: F45 < Y23, Y35 < Y21, F22 < Y10 < F4, F33. Indeed, NMR measurements and 1-ms-long simulation^3, 19^ observed residues F33 and F4 as fast flipping residues. We also investigated the correlation between solvent accessible surface area (SASA) of aromatic side-chains and experimental ring-flipping rates *k_exp_*, however, a lack of correlation was observed similar to the previous study^50^ (**figure S37** in SI). The free energy surface of solvent-exposed residues (F4, Y10) and buried residues (F22, F33, Y35) indicated multiple ring-flipping pathways. The circular correlation coefficient^51^ between *χ*_1_, *χ*_2_ dihedrals of eight aromatic residues obtained from 1-ms-long trajectory indicate no correlated motions between these residues (**figure S38 in SI**). The side-chain aromatic ring of residues Y23 and F45 flips around seconds; these time scales are still beyond the scope of present MD simulations. This study illustrates that InMetaD can be applied to understand motions occurring at second time scales, as also demonstrated by drug-unbinding studies^52–54^. The InMetaD simulations of residues F33 and F22 indicate friction effects during barrier crossing events. Further investigations is needed to understand the origin of these observations. This study illustrates the complexity of the ring flipping process in globular proteins.

## ASSOCIATED CONTENT

### Supporting Information

The Supporting Information is available free of charge at …

A convergence of WTMetaD simulations, results of *χ*_2_ InMetaD simulations, distribution of side-chain torsions of a C14-C38 disulfide bond, SASA results, circular cross-correlation of *χ*_2_ and *χ*_2_ dihedral values obtained from 1-ms-long D. E. Shaw Research (DESRES) trajectory, details of state-to-state transitions of residue F22, and details of VAC-metadynamics simulations.

## AUTHOR INFORMATION

### Author Contributions

The manuscript was written through the contributions of all authors. All authors have given approval to the final version of the manuscript.

### Funding Resources

PS acknowledges funding from the Swedish Research council (project 2017-05318). The computations were performed on resources provided by SNIC through LUNARC at Lund University and HPC2N at Umeå University under Projects SNIC 2019/3-317 and SNIC 2020/5-367.

### Notes

The authors declare no competing financial interest.

## ACKNOWLEDGMENT

M.K. thanks Dr. Yong Wang, Department of Biology, the University of Copenhagen, for providing bootstrap analysis Matlab script of transition timescales.

## For Table of Contents Only

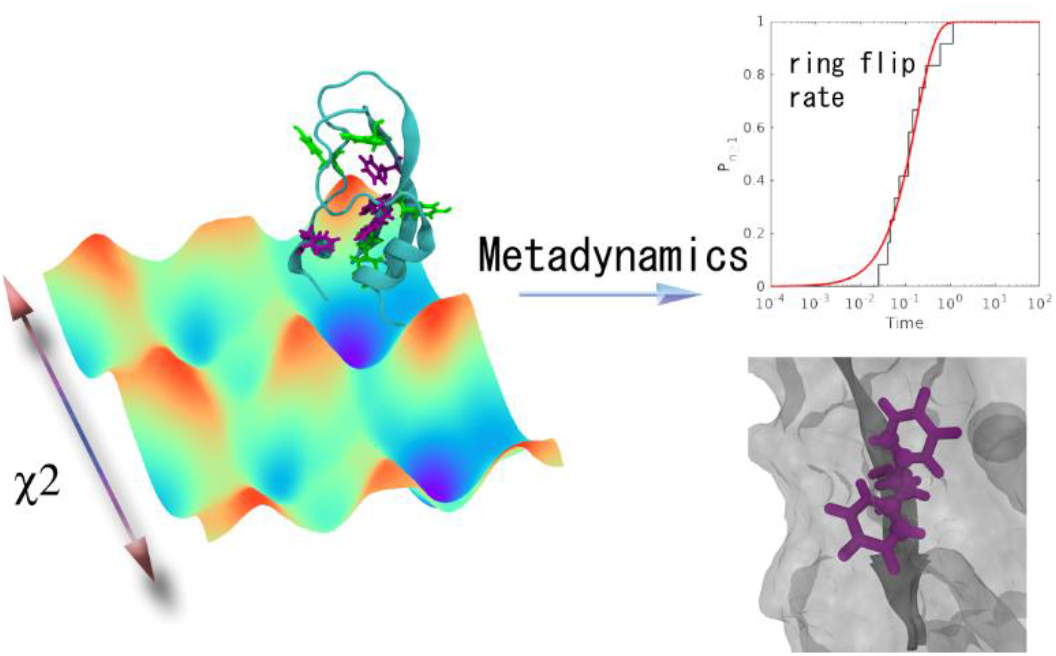

