## supplementary_material_v1 for "Free Energy Landscape and Rate Estimation of the Aromatic Ring Flips in Basic Pancreatic Trypsin Inhibitor Using Metadynamics"

**Table S1.** The values of $r_0$ and $d_0$ for $C_\alpha$ atom pairs in CMAP (Equation 2)

| | $C_\alpha$ atom pair (***ij***) | | | | **Figure with atom names** |
|---|---|---|---|---|---|
| Sr. no. | $C_\alpha(i)$ | $C_\alpha(j)$ | $r_0$ (nm) | $d_0$ (nm) | 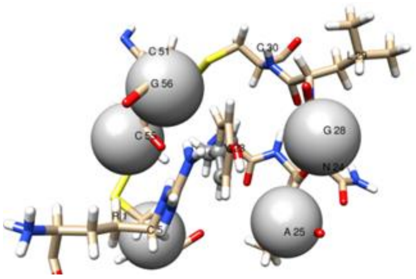 |
| 1 | A25 | G28 | 0.08 | 0.47 | |
| 2 | G28 | G56 | 0.10 | 0.77 | |
| 3 | C55 | A25 | 0.08 | 0.92 | |
| 4 | C55 | C5 | 0.03 | 0.58 | |
| 5 | A25 | C5 | 0.06 | 0.72 | |

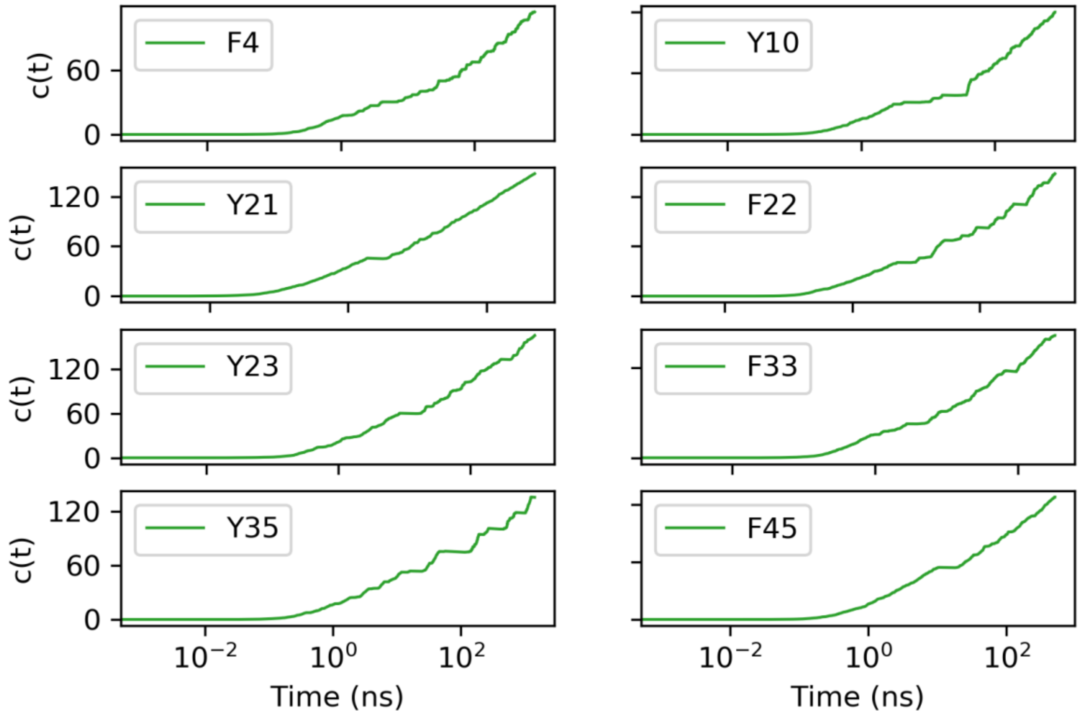


**Figure S1.** The bias offset c(t) obtained from reweighting converged simulations for each residue. Please note, residue Y35 $\chi_2 - \chi_1$ metadynamics simulation lack convergence.

**Simulation results:**

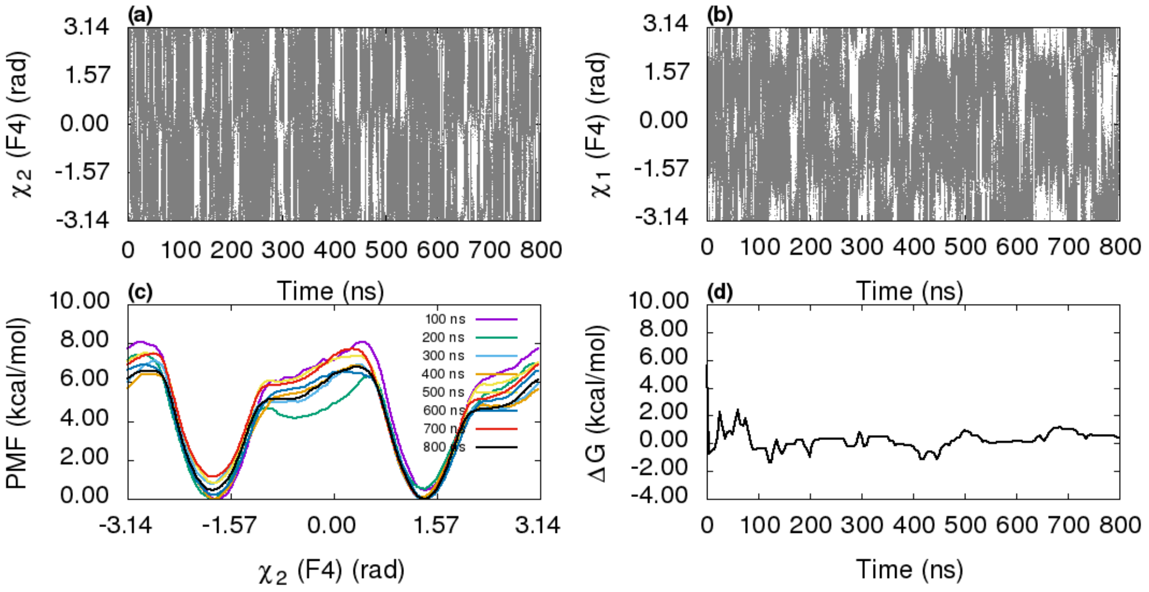


**Figure S2.** Diffusion of $\chi_2$ and $\chi_1$ during $\chi_2 - \chi_1$ metadynamics simulation of residue F4 is shown in figures (a) and (b), respectively. (c) Projection of free energy on $\chi_2$ in time intervals of 100 ns. (d) The free energy difference between two symmetric states along $\chi_2$.

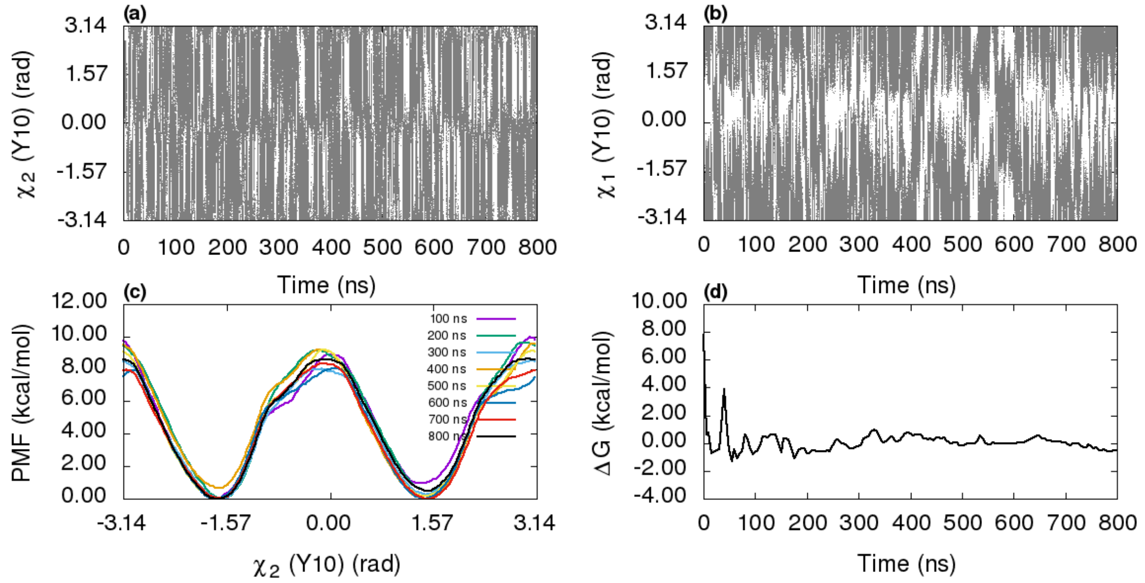


**Figure S3.** The time series of $\chi_2$ and $\chi_1$ during $\chi_2 - \chi_1$ metadynamics simulation showed in figures (a) and (b), respectively. (c) the projection of free energy on $\chi_2$ in time intervals of 100 ns. (d) The free energy difference between two symmetric states along $\chi_2$.

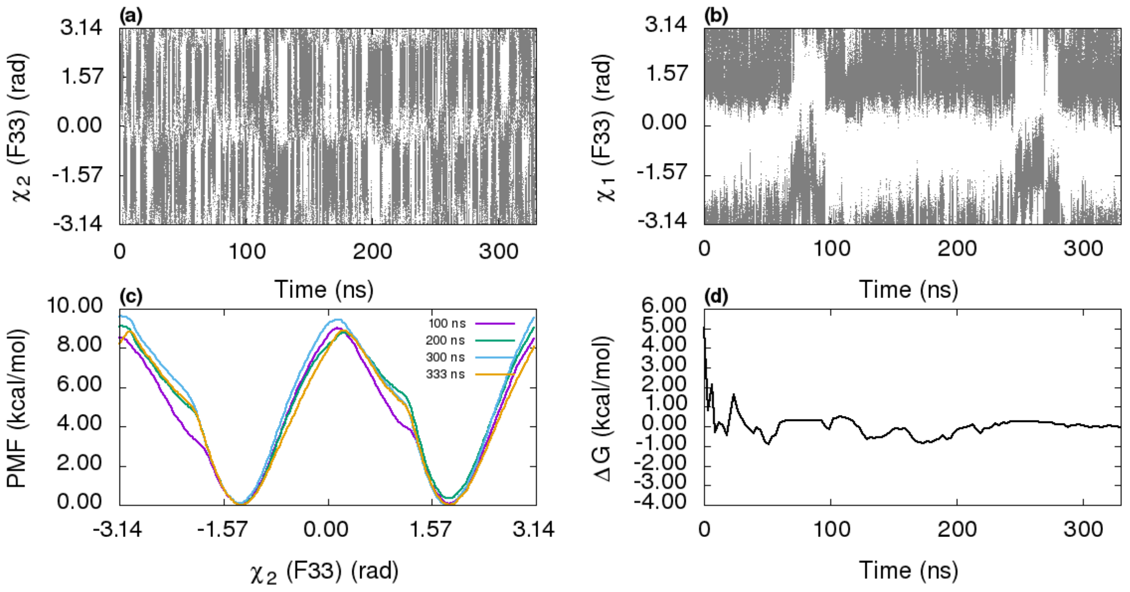


**Figure S4.** The diffusion of $\chi_2$ and $\chi_1$ variables during $\chi_2 - \chi_1$ metadynamics simulation of residue F33 (figures (a) and (b), respectively). (c) the projection of free energy on $\chi_2$ in time intervals of 100 ns. (d) The free energy difference between two symmetric states along $\chi_2$.

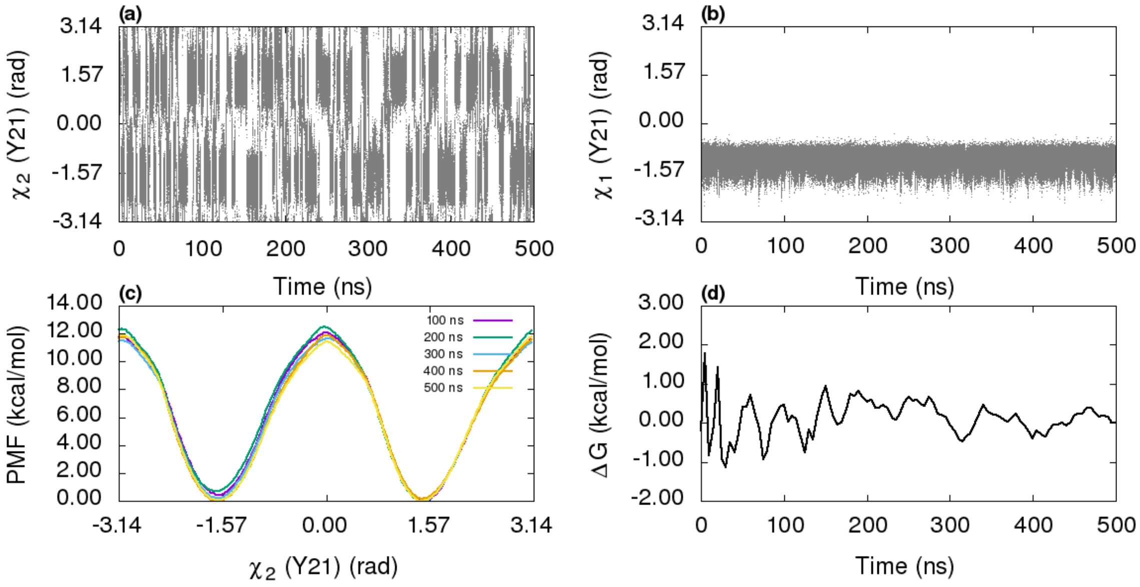


**Figure S5.** The diffusion of $\chi_2$ and $\chi_1$ collective variables during $\chi_2$ metadynamics simulation of residue Y21 (figures (a) and (b), respectively). (c) the free energy along

$\chi_2$ in the time intervals of 100 ns. (d) The free energy difference between two symmetric states along $\chi_2$.

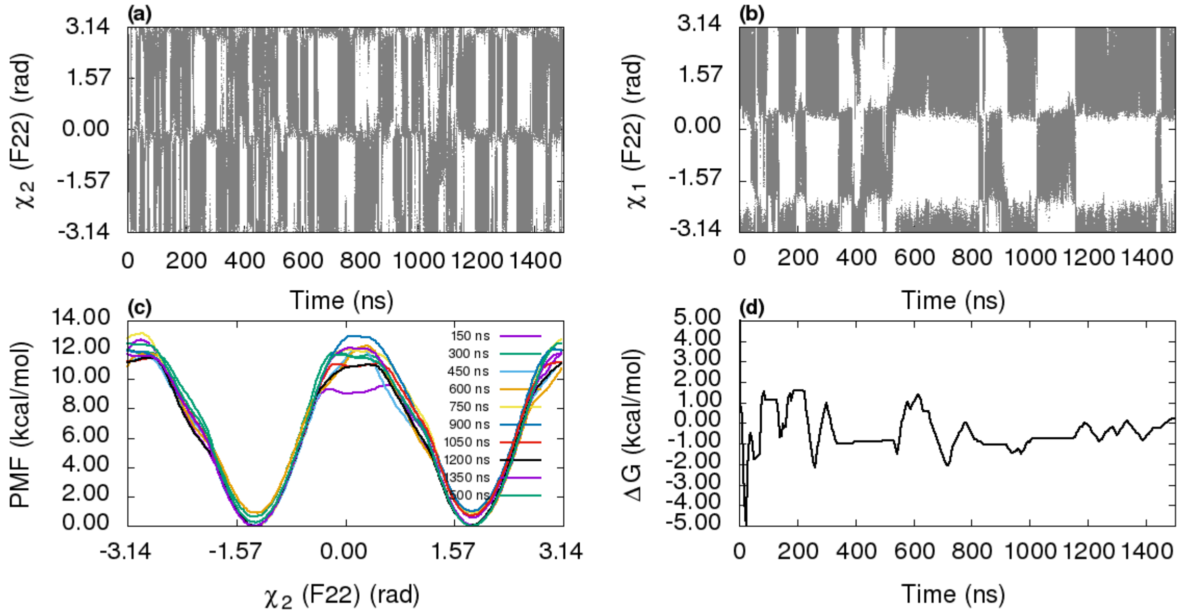


**Figure S6.** The time series of $\chi_2$ and $\chi_1$ during $\chi_2 - \chi_1$ metadynamics simulation of residue F22 showed in figures (a) and (b), respectively. (c) the projection of free energy on $\chi_2$ in time intervals of 150 ns. (d) The free energy difference between two symmetric states along $\chi_2$.

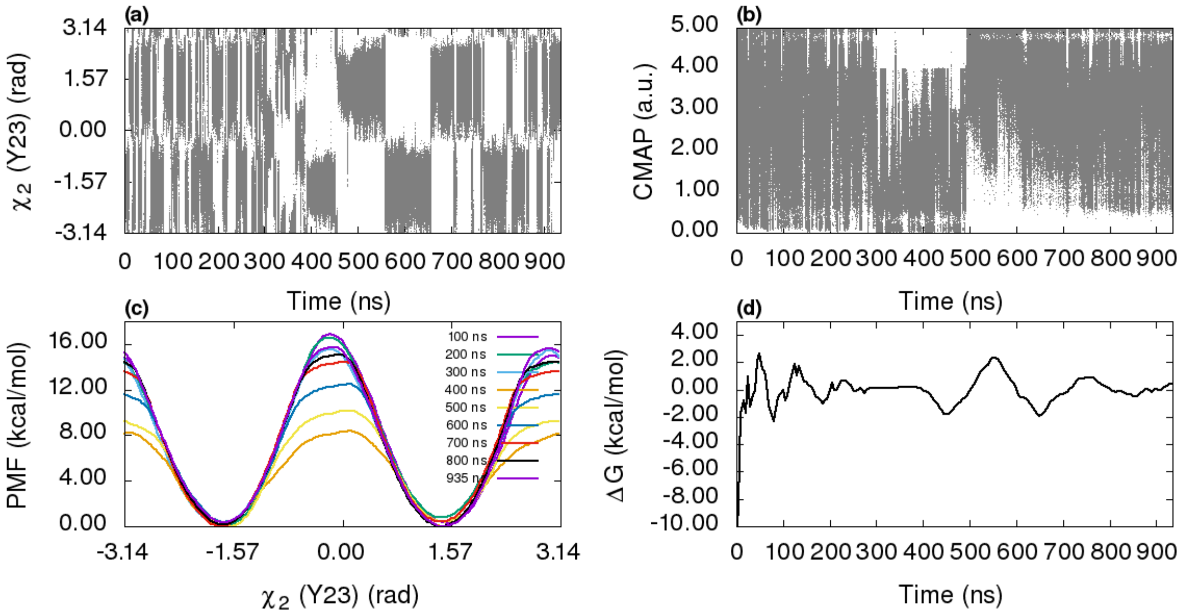


**Figure S7.** The time series of $\chi_2$ and contact map (CMAP) collective variables during $\chi_2$-CMAP metadynamics simulation of residue Y23 in figures (a) and (b), respectively. (c) the projection of free energy along $\chi_2$ in time intervals of 100 ns. (d) the free energy difference between two symmetric states along $\chi_2$.

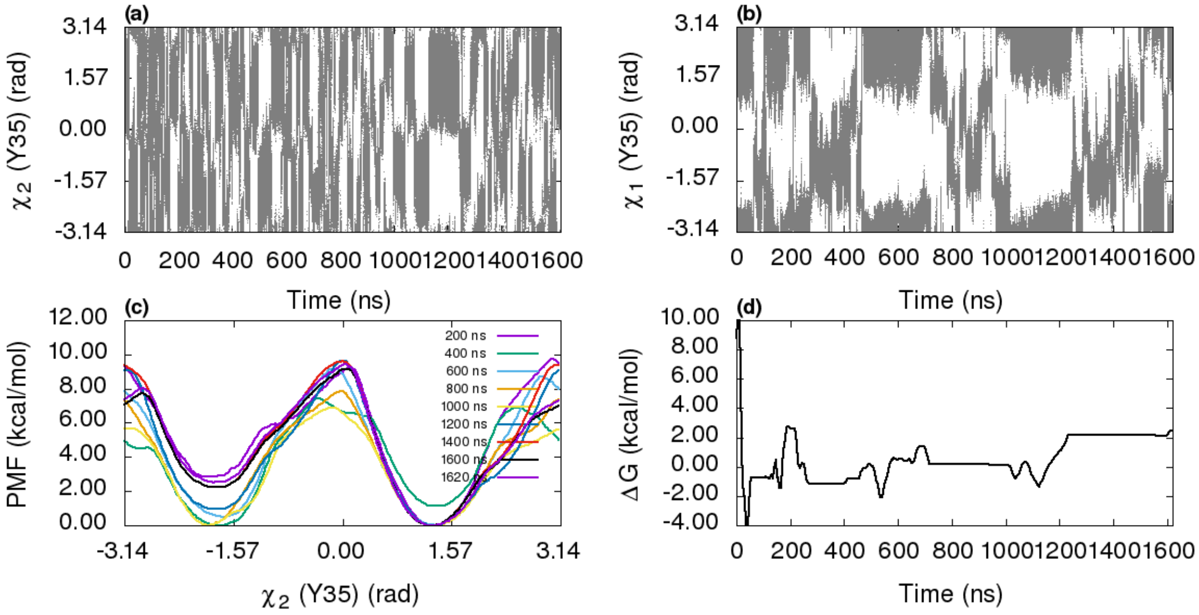


**Figure S8.** The time series of $\chi_2$ and $\chi_1$ during $\chi_2 - \chi_1$ metadynamics simulation of residue Y35 in figures (a) and (b), respectively. (c) the projection of free energy on $\chi_2$

in time intervals of 200 ns. (d) The free energy difference between two symmetric states along $\chi_2$.

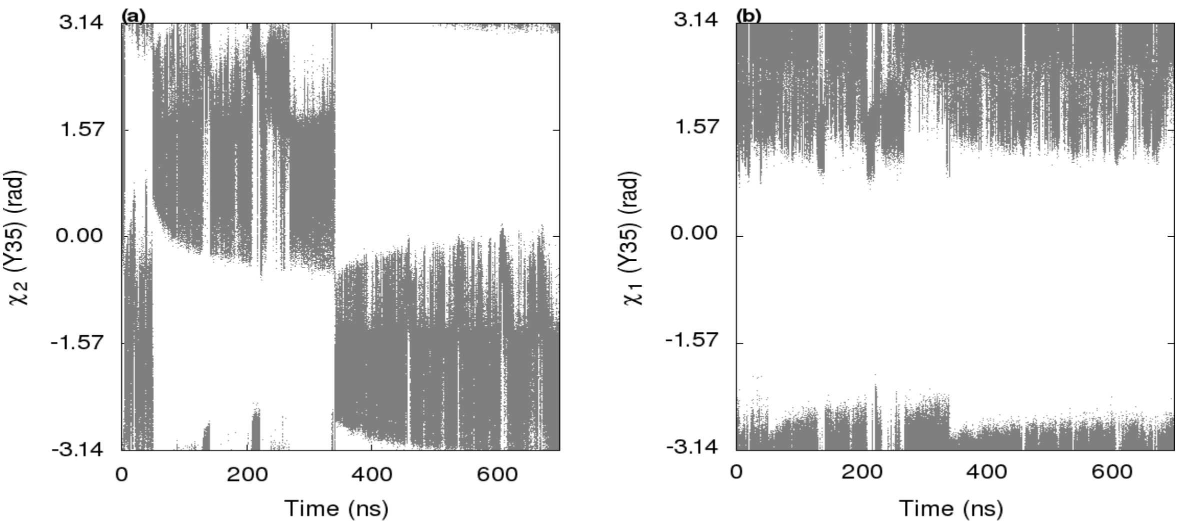


**Figure S9.** The time series of $\chi_2$ and $\chi_1$ during $\chi_2$ metadynamics simulation of residue Y35 in figures (a) and (b), respectively.

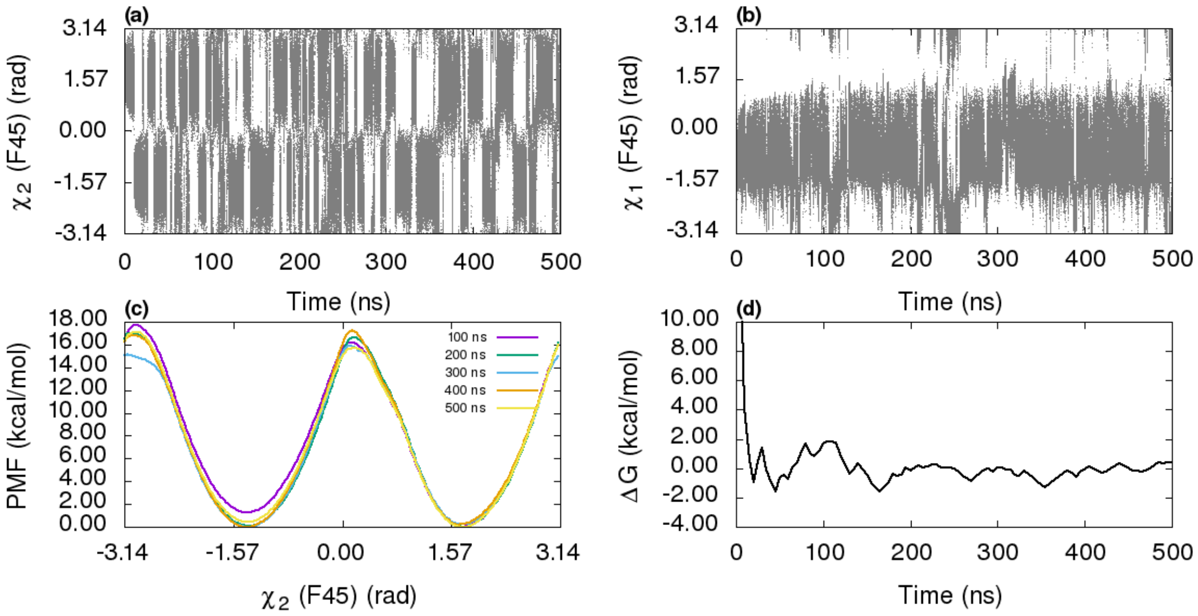


**Figure S10.** The diffusion of $\chi_2$ and $\chi_1$ during $\chi_2 - \chi_1$ metadynamics simulation of residue F45 in figures (a) and (b), respectively. (c) the projection of free energy on $\chi_2$

in time intervals of 100 ns. (d) The free energy difference between two symmetric states along $\chi_2$.

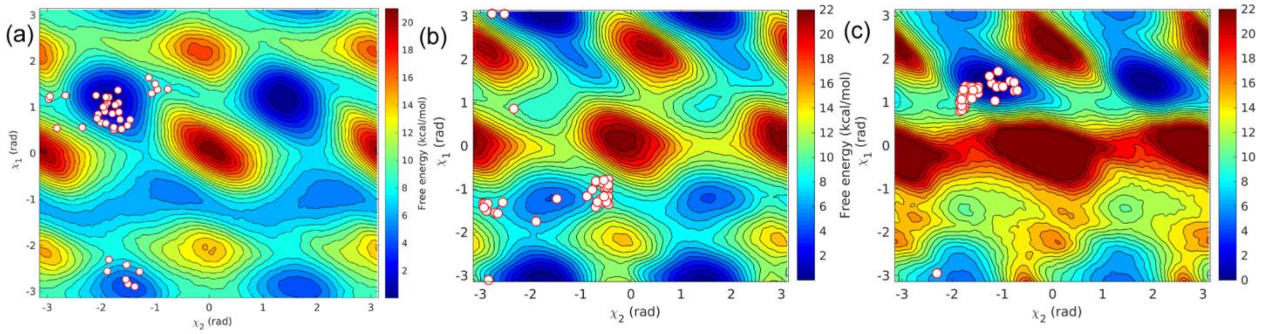


**Figure S11.** The distribution of trajectory endpoints after flipping for (a) residue F4, (b) residue Y10, and (c) residue F33 ($\chi_2 - \chi_1$ metadynamics simulations).

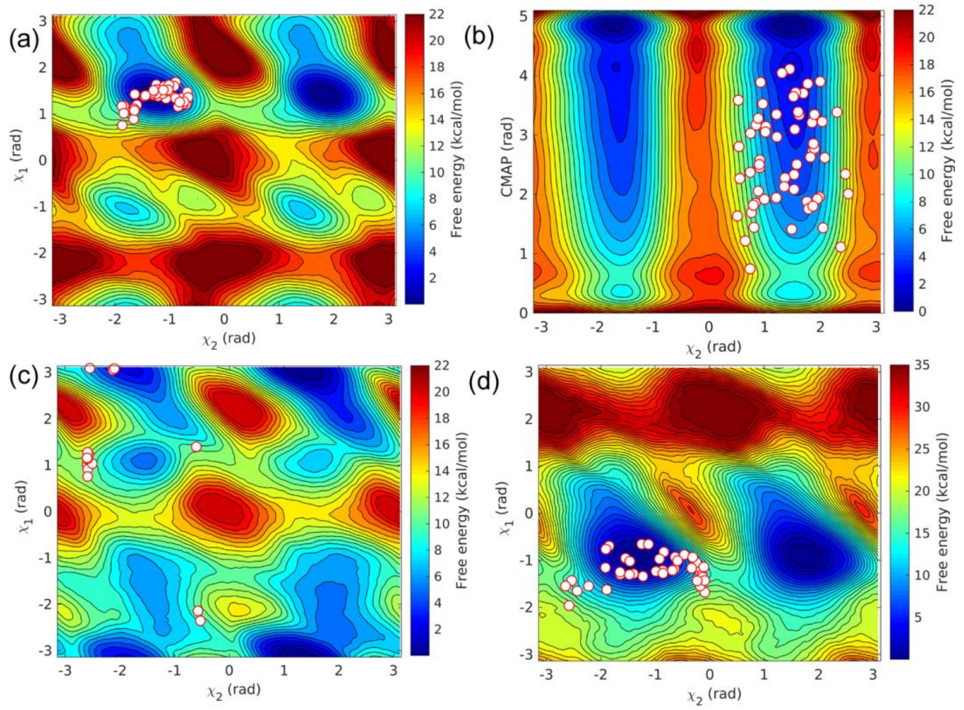


**Figure S12.** The distribution of trajectory endpoints after flipping for (a) residue F22, (b) residue Y23, (c) residue Y35, and (d) residue F45.

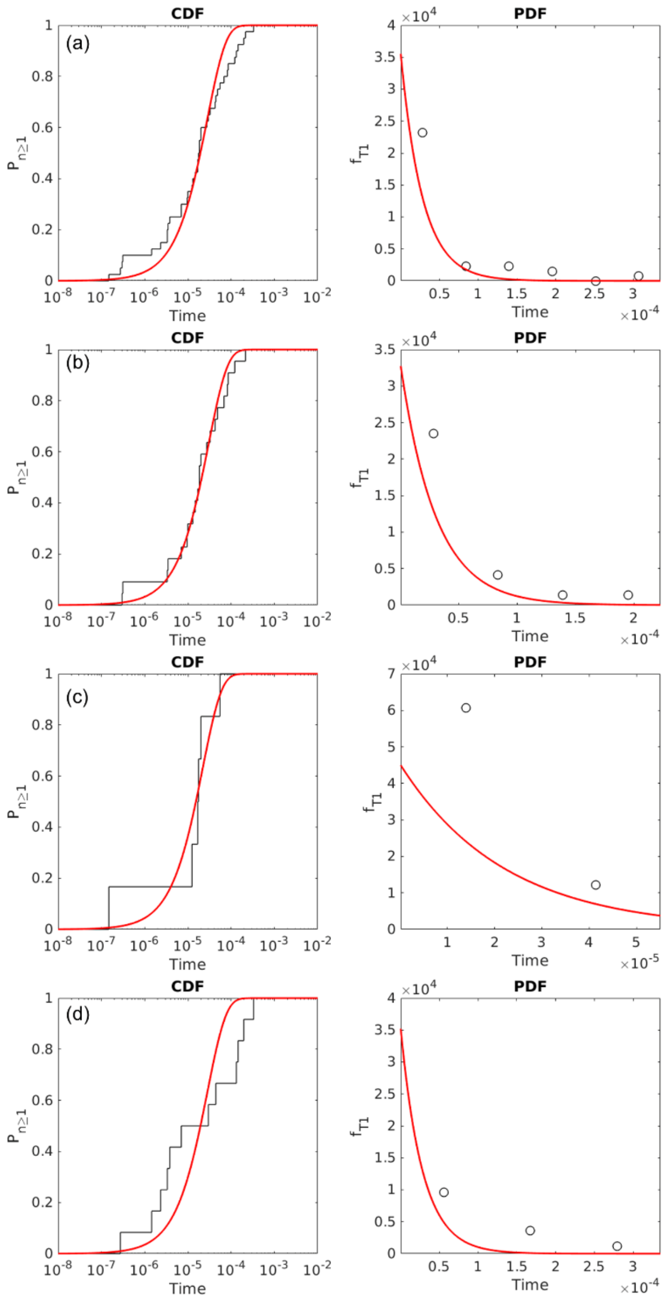


**Figure S13**. Theoretical Poisson fit to cumulative distribution of transition timescales obtained from $\chi_2 - \chi_1$ infrequent metadynamics simulations of residue F4 by considering (a) all 40 simulations (p-value = 0.32 and μ ln 2/tm = 1.79), (b) 22 $\chi_2 - \chi_1$assisted A-I1-I1' transition (p-value = 0.82 and μ ln 2/tm = 1.48) , (c) 6 A-I2' transitions (p-value = 0.71 and μ ln 2/tm = 0.80) , (d) 12 A-A' direct transitions (p-value = 0.13 and μ ln 2/tm = 2.78).

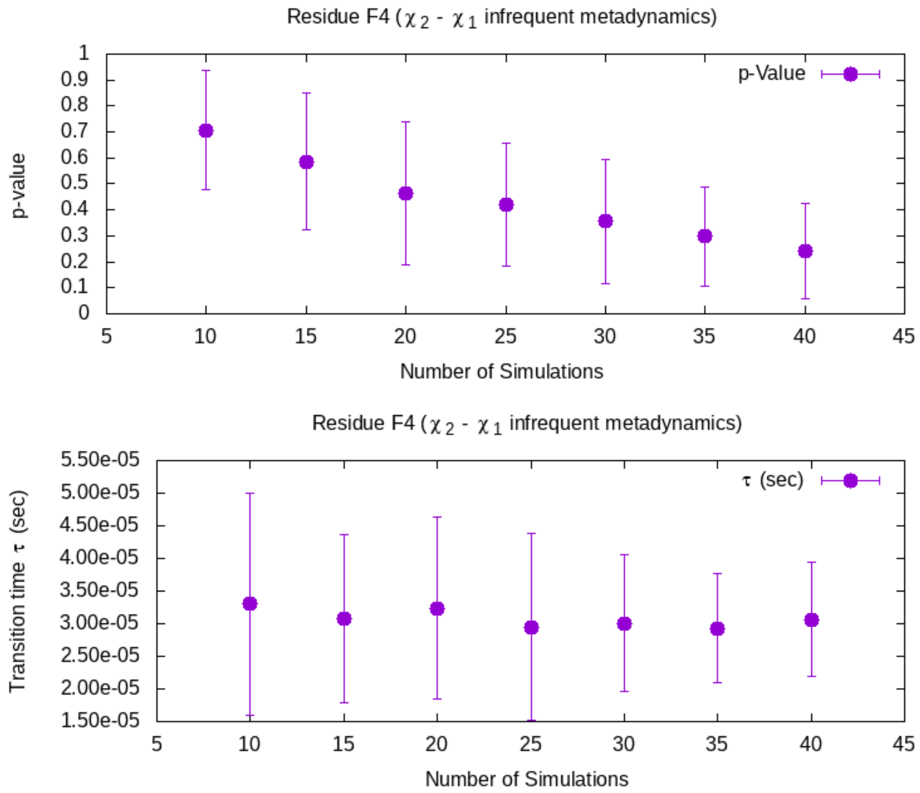


**Figure S14.** Errors in p-value and transition timescales from $\chi_2 - \chi_1$ infrequent metadynamics simulations of residue F4 obtained with bootstrapping.

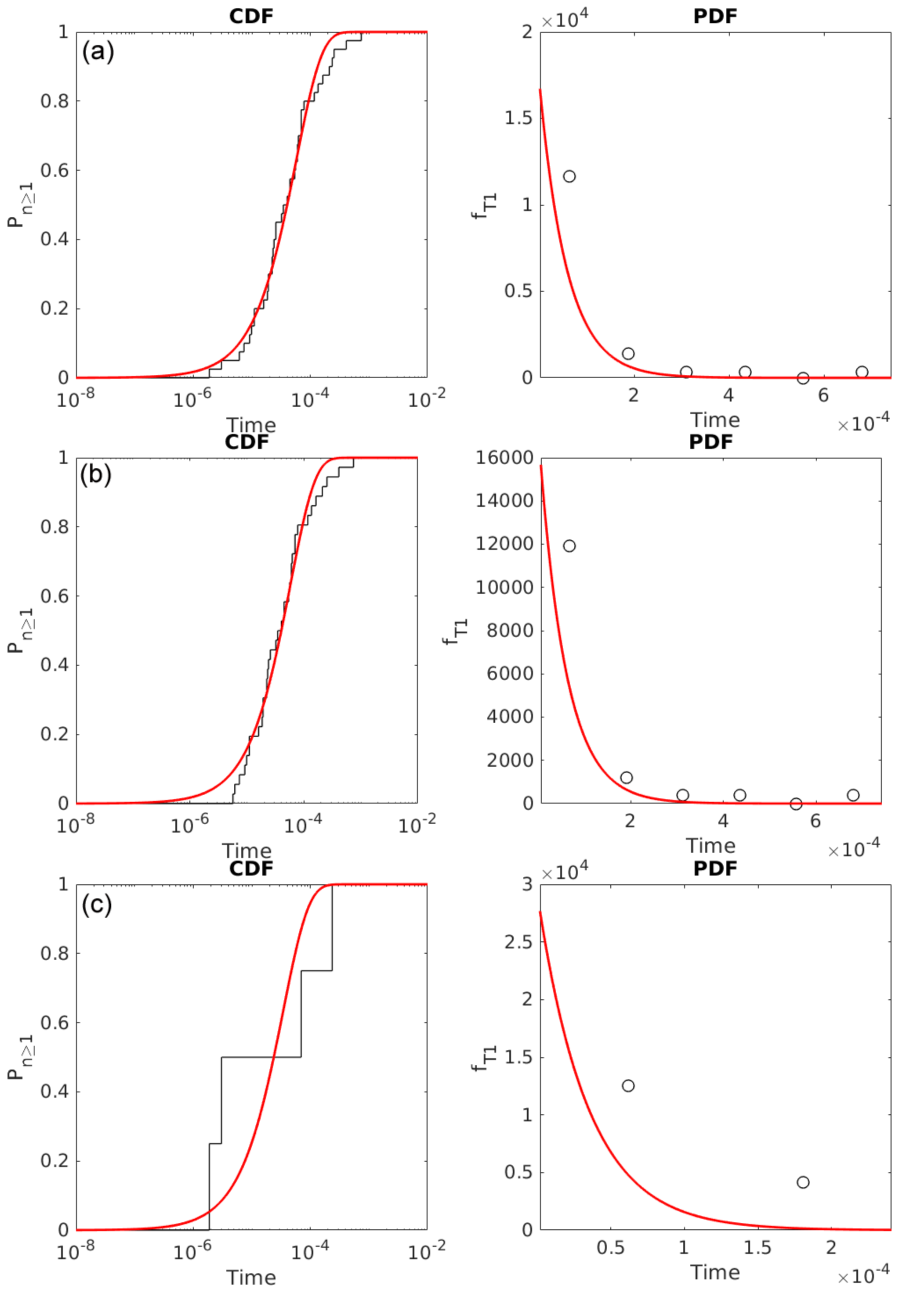


**Figure S15.** Theoretical Poisson fit to cumulative distribution of transition timescales obtained from χ2- χ1 infrequent metadynamics simulations of residue Y10 by considering (a) all 40 simulations (p-value = 0.80 and μ ln 2/tm = 1.54), (b) 36 $\chi_2 - \chi_1$ driven A-I1-I1' transition (p-value = 0.89 and μ ln 2/tm = 1.54), (c) 4 A-A' direct transitions (p-value = 0.39 and μ ln 2/tm = 1.48).

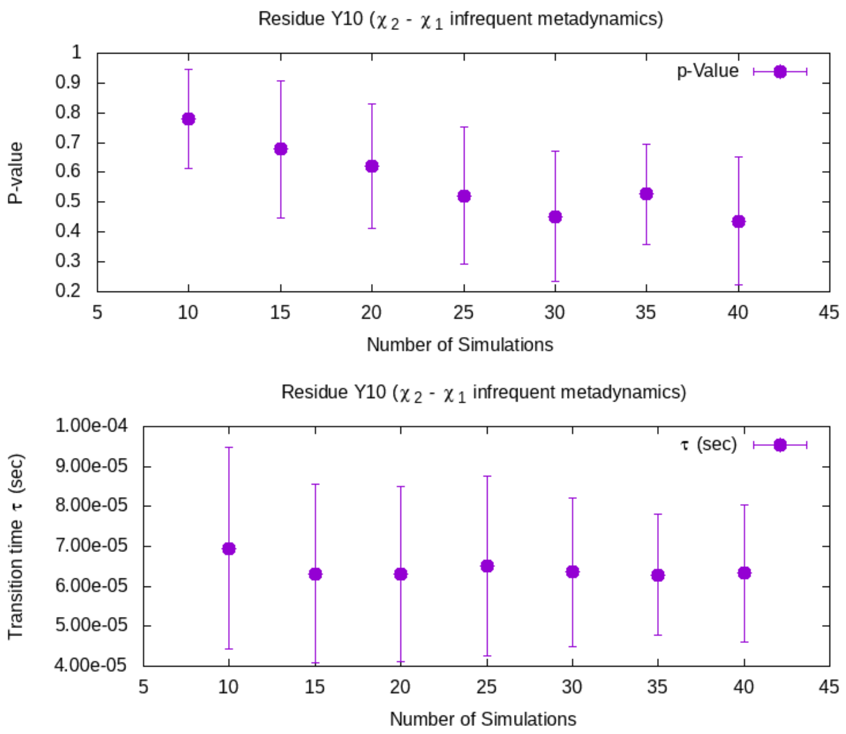


**Figure S16.** Errors in p-value and transition timescales from $\chi_2 - \chi_1$infrequent metadynamics simulations of residue Y10 obtained with bootstrapping.

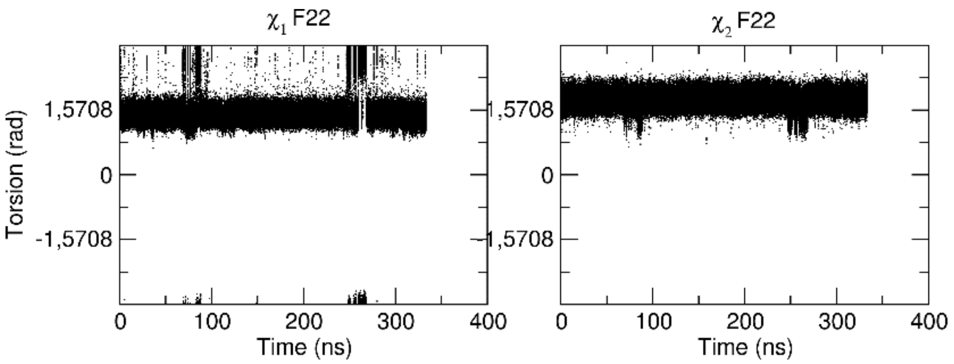


**Figure S17.** $\chi_1$ and $\chi_2$ torsion values for residue F22 during metadynamics simulation of residue F33.

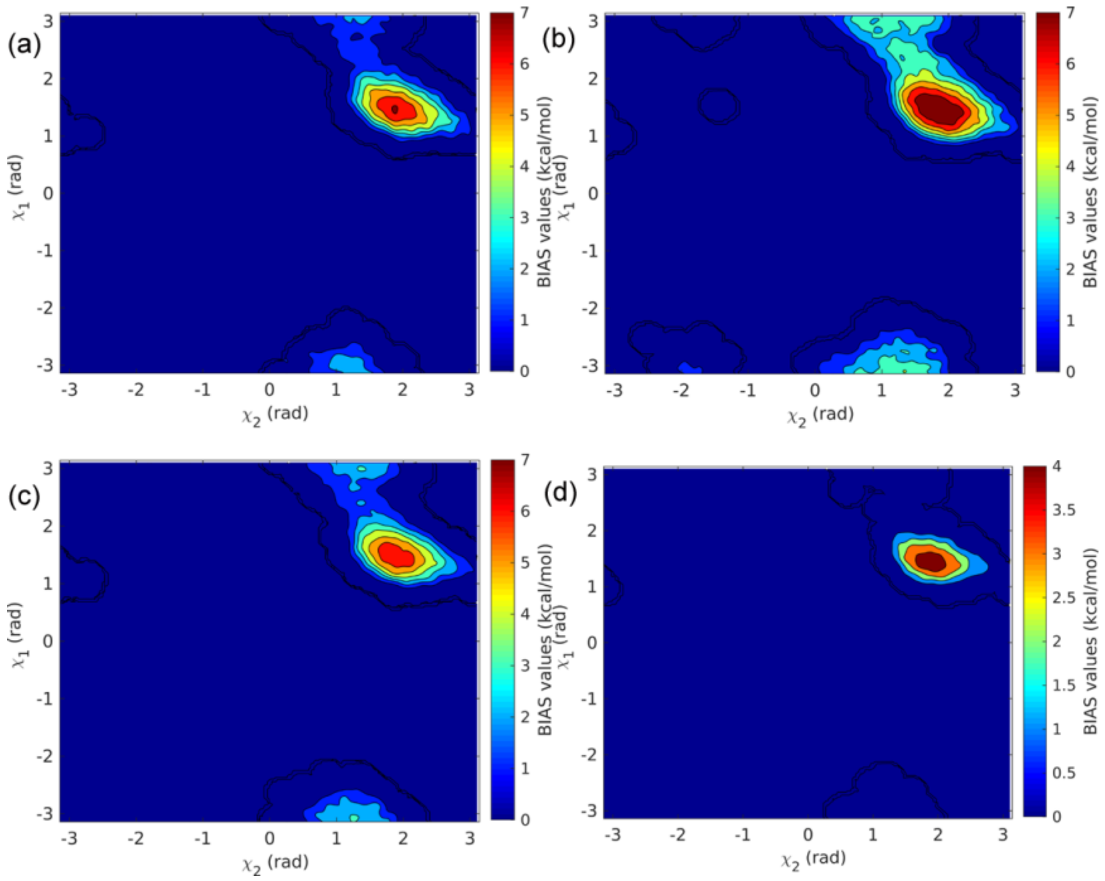


**Figure S18.** The illustration of bias deposition in the barrier region along $\chi_1$ during infrequent metadynamics simulation along $\chi_2 - \chi_1$ of F33 in some of the trajectories in panels (a)-(c). Panel (d) shows an example where bias is not deposited in the TS region.

**Table S2.** Results of infrequent metadynamics simulations of residue F33 with wall bias and without wall bias.

| Residue F33 | Number of simulations | $t_{flip}$ (s) | $k_{flip}$ $(s^{-1})$ | p-value | μ ln 2/tm |
|---|---|---|---|---|---|
| No wall bias | 40 | $3.50 \times 10^{-5}$ | $2.86 \times 10^{4}$ | 0.03 | 2.47 |
| With wall bias | 12 | $3.14 \times 10^{-5}$ | $3.18 \times 10^{4}$ | 0.52 | 3.78 |

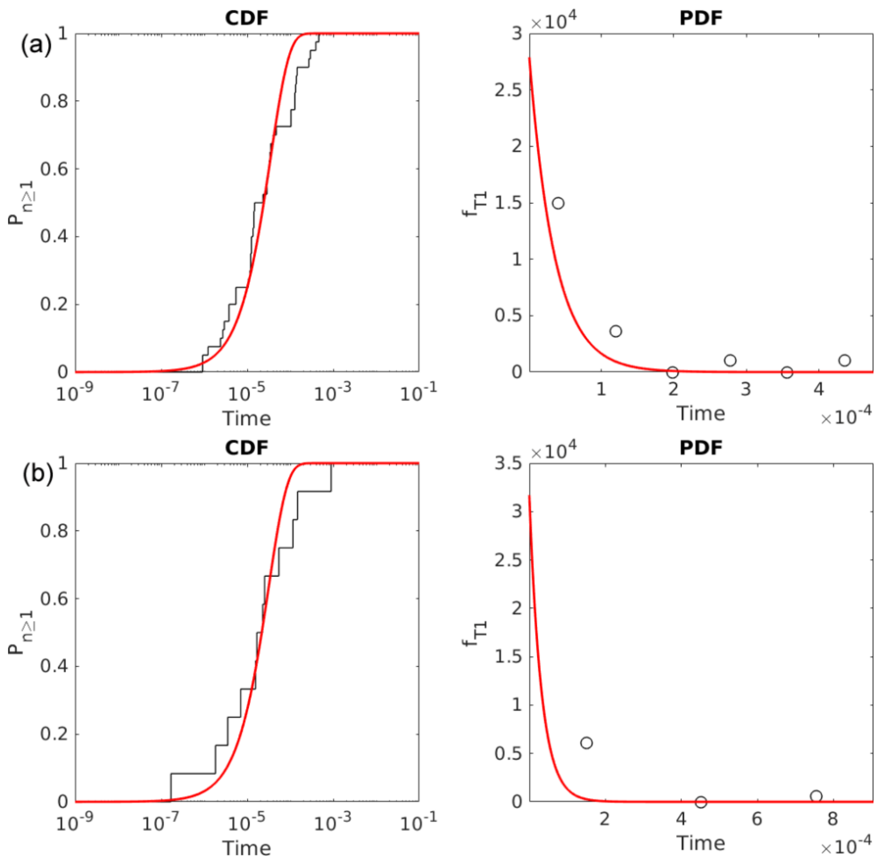


**Figure S19.** Theoretical Poisson fit to the cumulative distribution of transition timescales obtained from $\chi_2 - \chi_1$ infrequent metadynamics simulations of residue F33

(a) without repulsive walls, and (b) with repulsive wall potential (UPPER_WALL) at $\chi 1$ = 2.4 rad to avoid bias deposition on transition state (12 simulations).

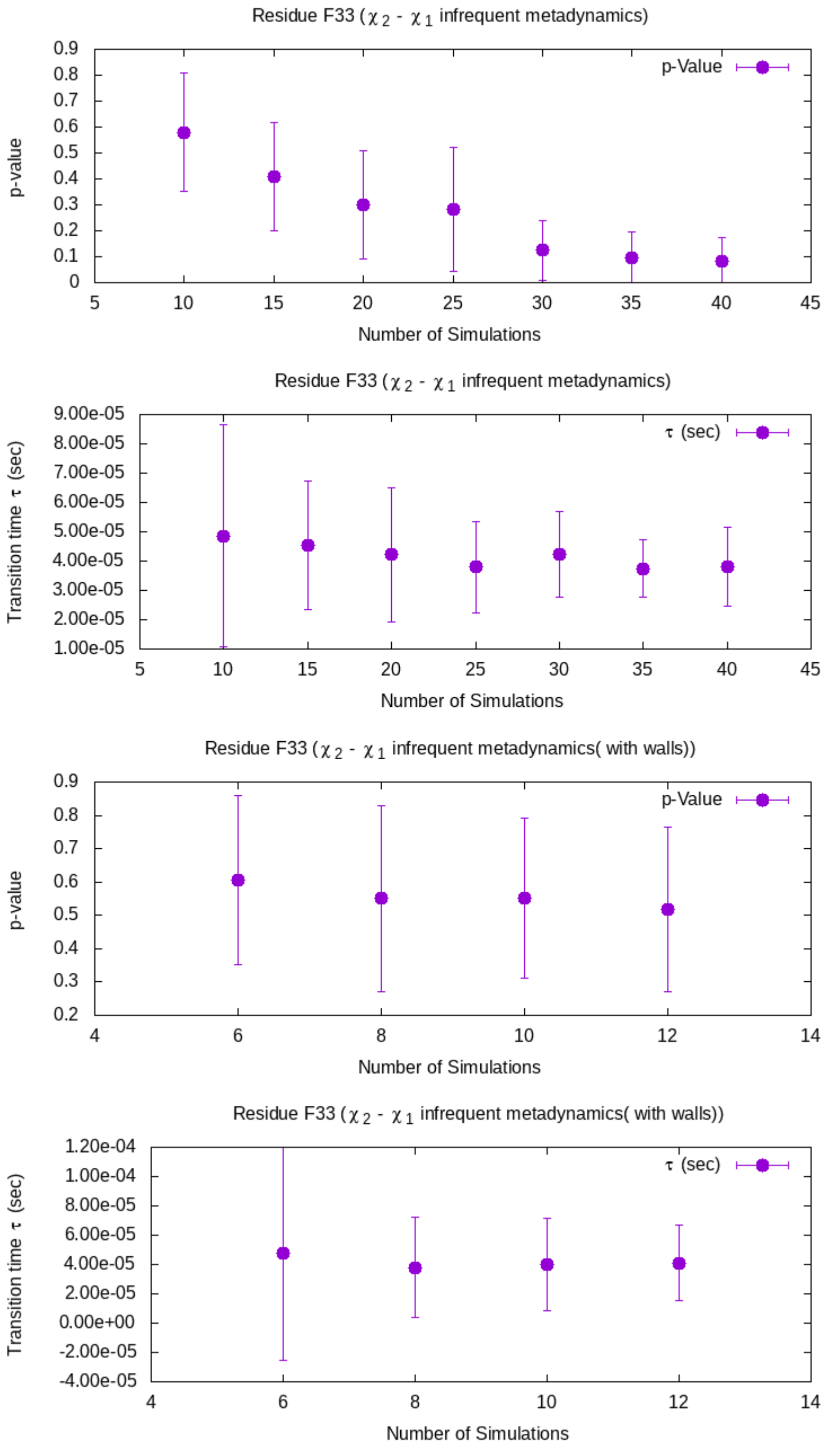

**Figure S20.** Errors in p-value and transition timescales from $\chi_2 - \chi_1$ infrequent metadynamics simulations of residue F33 without walls (upper two panels) and with wall bias (lower two panels) obtained with bootstrapping.

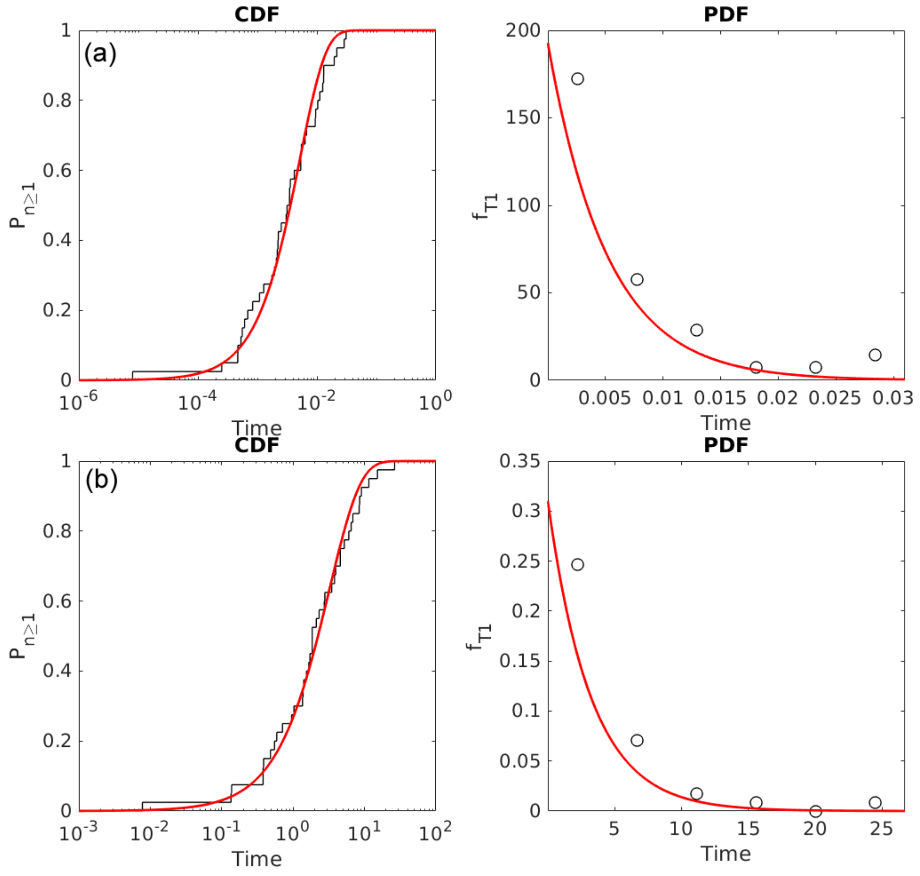


**Figure S21.** Theoretical Poisson fit to the cumulative distribution of transition timescales obtained from $\chi_2 - \chi_1$ infrequent metadynamics simulations of (a) residue Y21 and (b) F45.

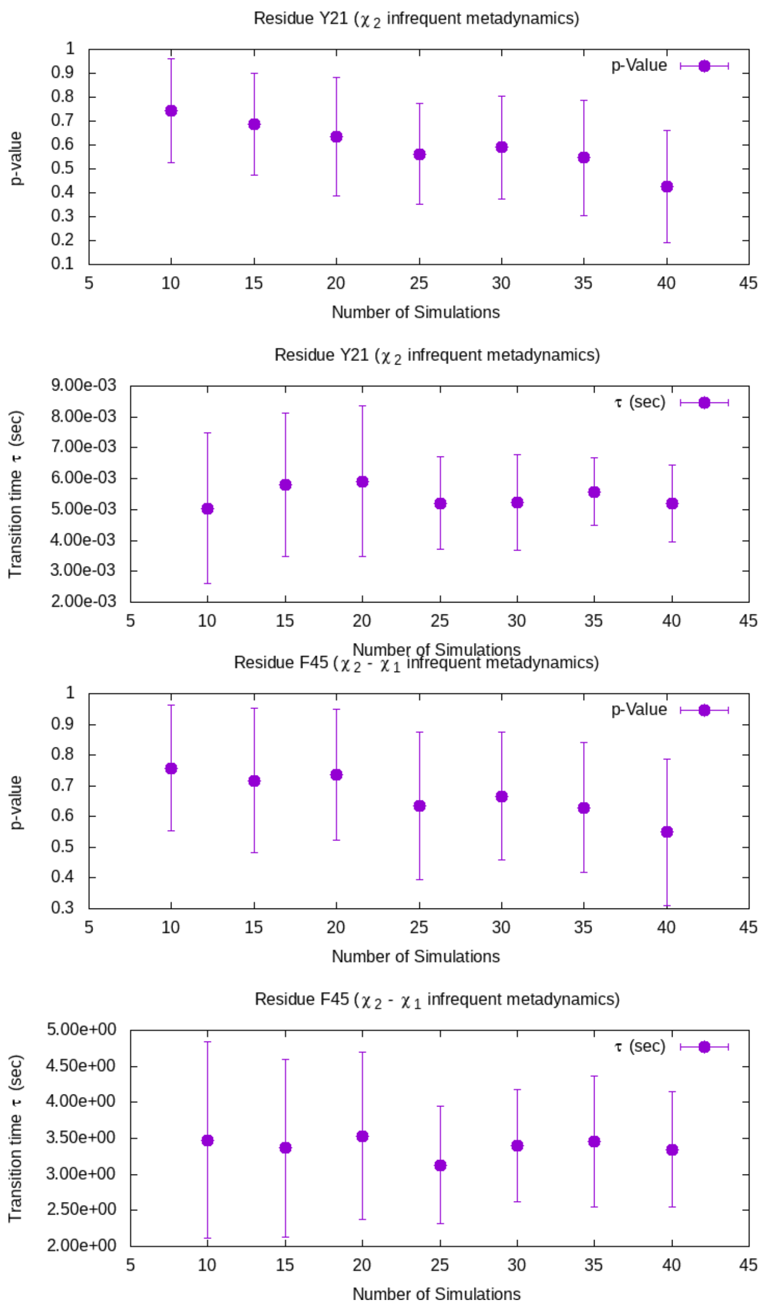


**Figure S22.** Errors in p-value and transition timescales from $\chi_2$ infrequent metadynamics simulations of residue Y21 (upper two panels) and $\chi_2 - \chi_1$biased residue F45 (lower two panels) obtained with bootstrapping.

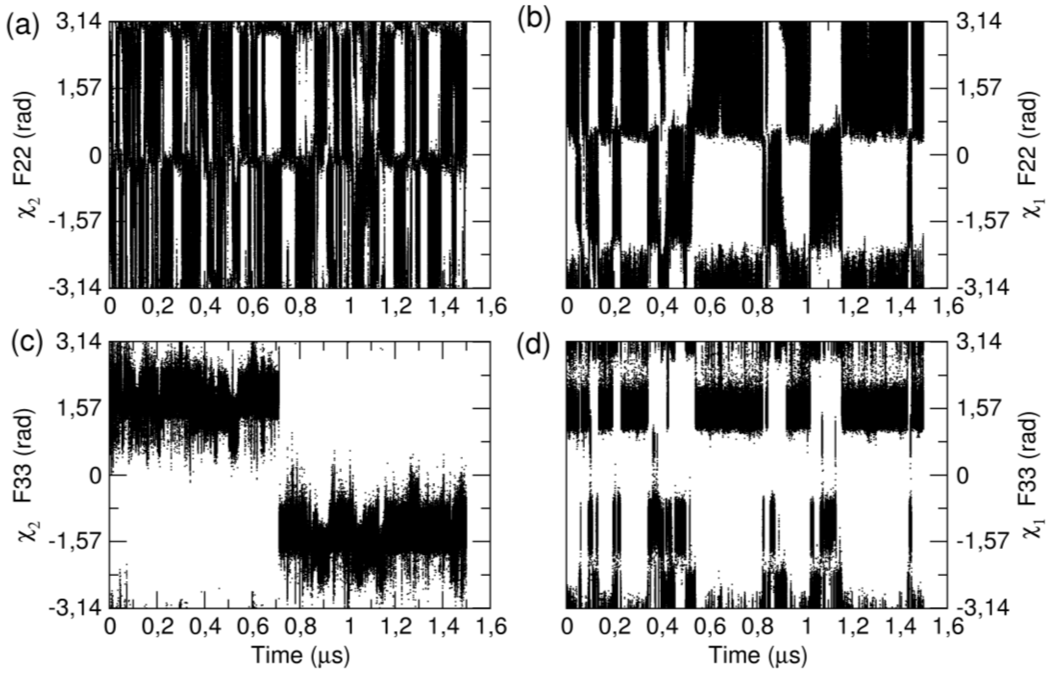


**Figure S23.** $\chi_1$ and $\chi_2$ torsion values for residue F22 and F33 in the metadynamics simulation of residue F22.

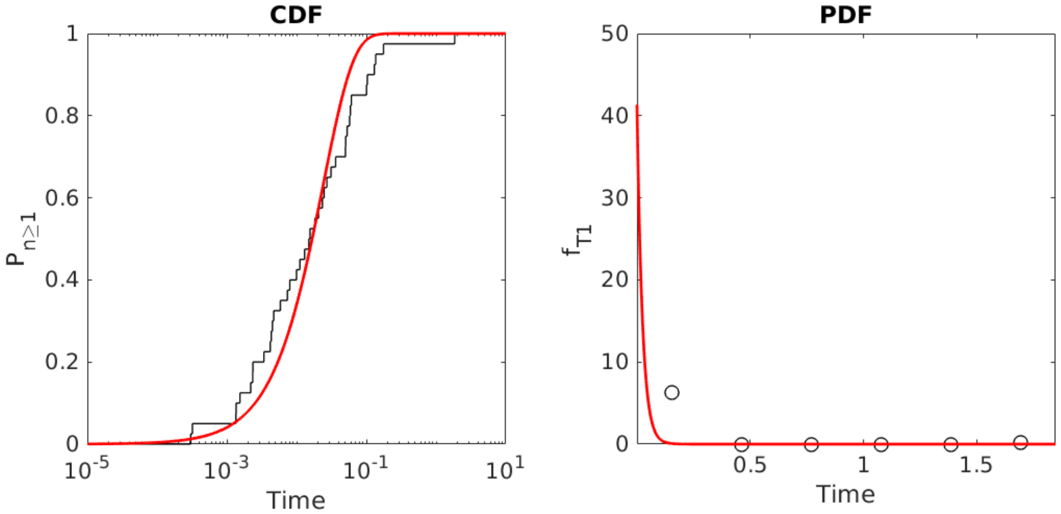


**Figure S24.** Theoretical Poisson fit to the cumulative distribution of transition timescales obtained from $\chi_2 - \chi_1$ infrequent metadynamics simulations of residue F22.

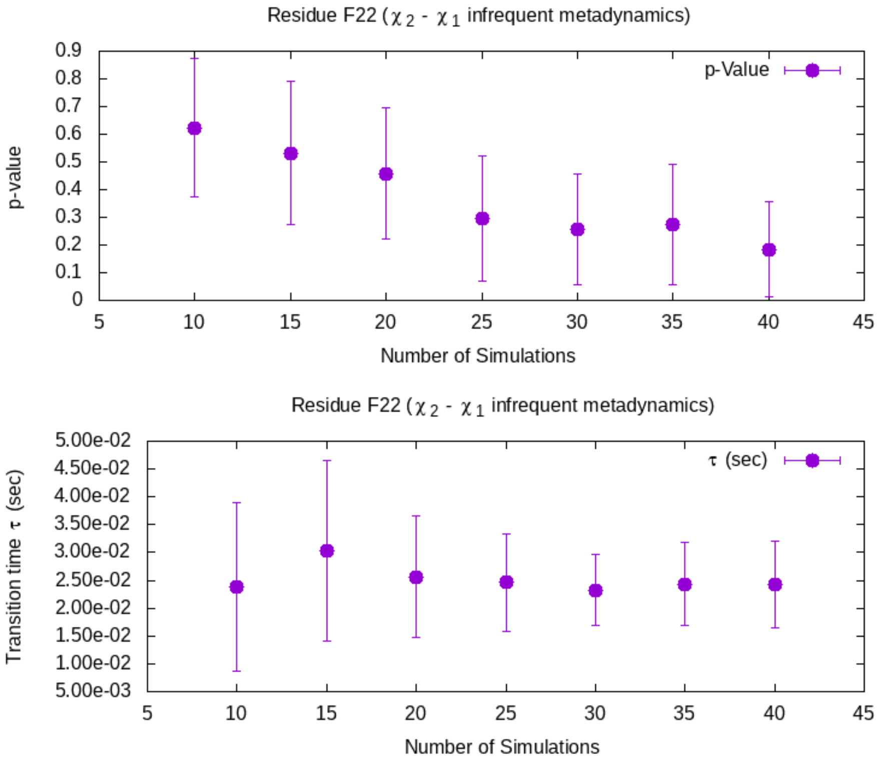


**Figure S25.** Errors in p-value and transition timescales from $\chi_2 - \chi_1$ infrequent metadynamics simulations of residue F22.

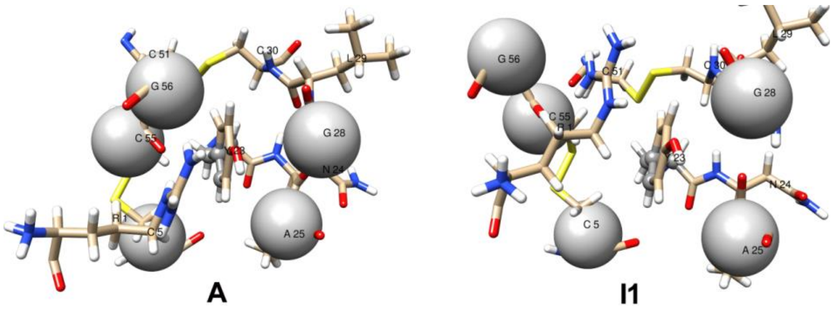


**Figure S26.** Representative structures observed in each minimum of $\chi_2$-CMAP free energy surface of residue Y23.

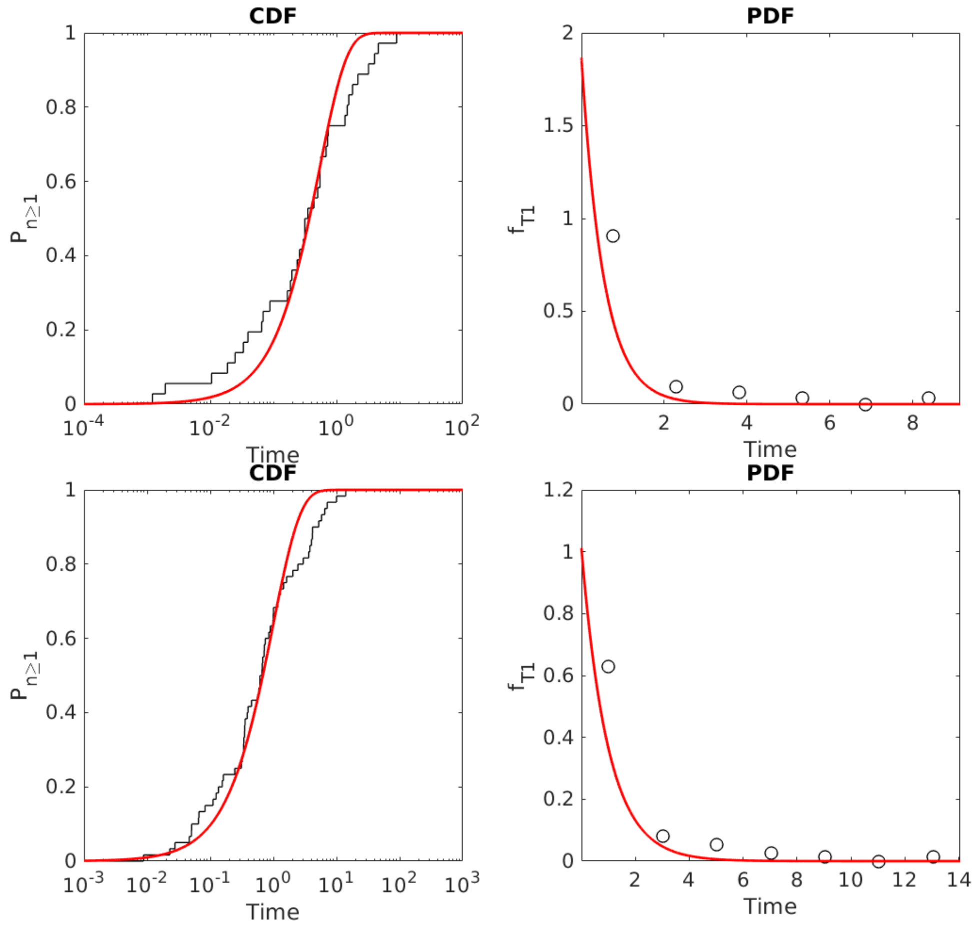


**Figure S27.** Theoretical Poisson fit to the cumulative distribution of $\tau_i\ (s)$ obtained from $\chi_2 - \chi_1$ (top panel, 26 simulations, p-value = 0.22 and μ ln 2/tm = 2.17) and $\chi_2$-CMAP (bottom panel, 60 simulations, p-value = 0.08 and μ ln 2/tm = 1.80) InMetaD simulations of residue Y23.

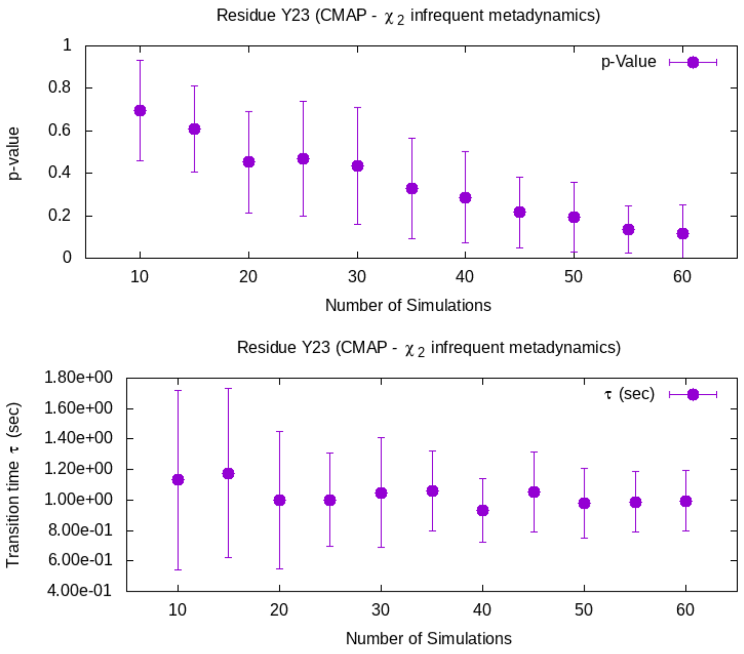


**Figure S28.** Errors in p-value and transition timescales from $\chi_2$- CMAP biased infrequent metadynamics simulations of residue Y23.

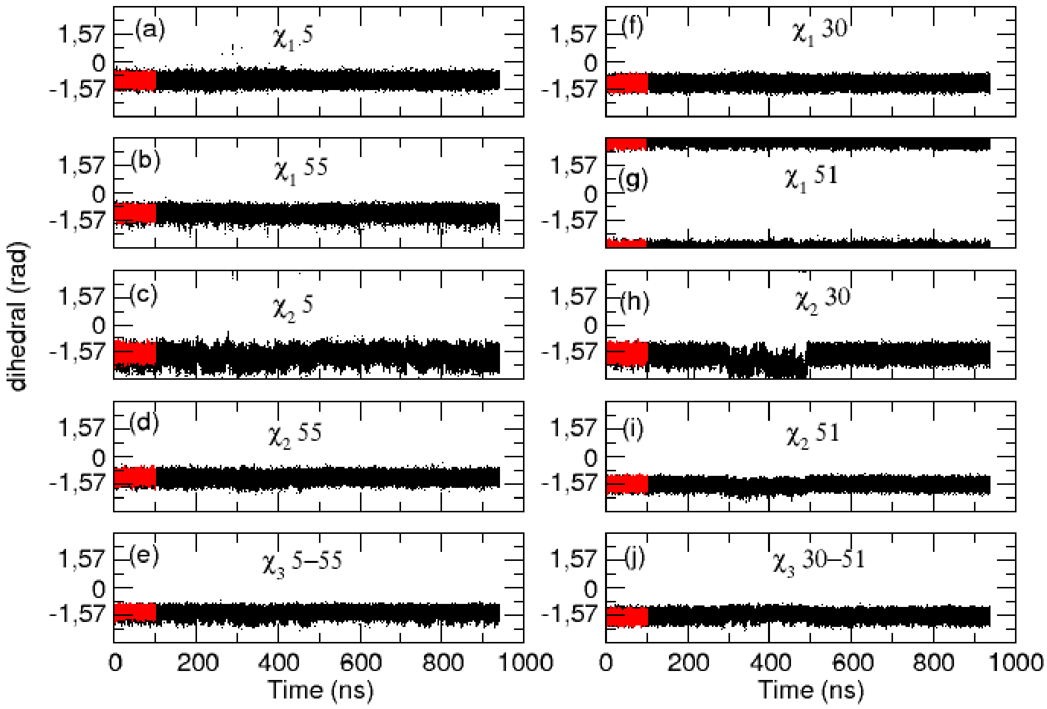


**Figure S29.** Values of $\chi_2$, $\chi_1$ , and $\chi_3$ torsions of 5-55 and 30-51 disulfide bridges from $\chi_2$-CMAP metadynamics simulation of residue Y23 (black) and 100 ns unbiased simulation (red).

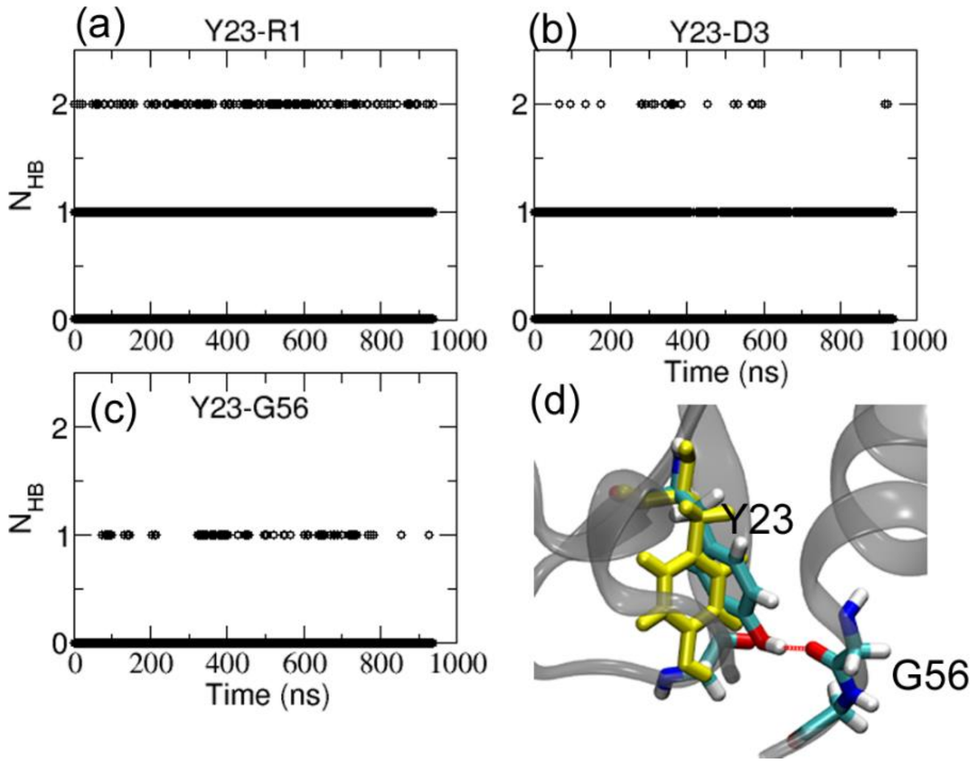


**Figure S30.** The number of hydrogen bonds between residues (a) Y23-R1, (b) Y23-D3, (c) Y23-G56, and (d) representative structure showing the hydrogen bonding of Y23 with G56 during the ring flip. The native orientation of the Y23 aromatic side-chain before flipping is shown in yellow color.

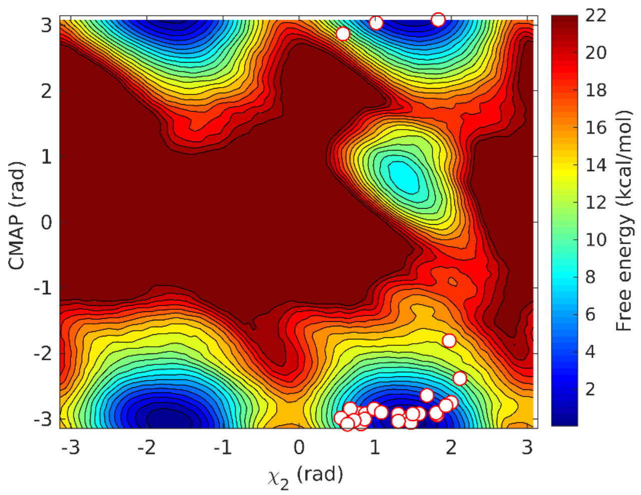


**Figure S31.** Free energy landscape of $\chi_2 - \chi_1$ biased metadynamics simulation of residue Y23. White circles indicate the 26 trajectory endpoints immediate after the ring flip.

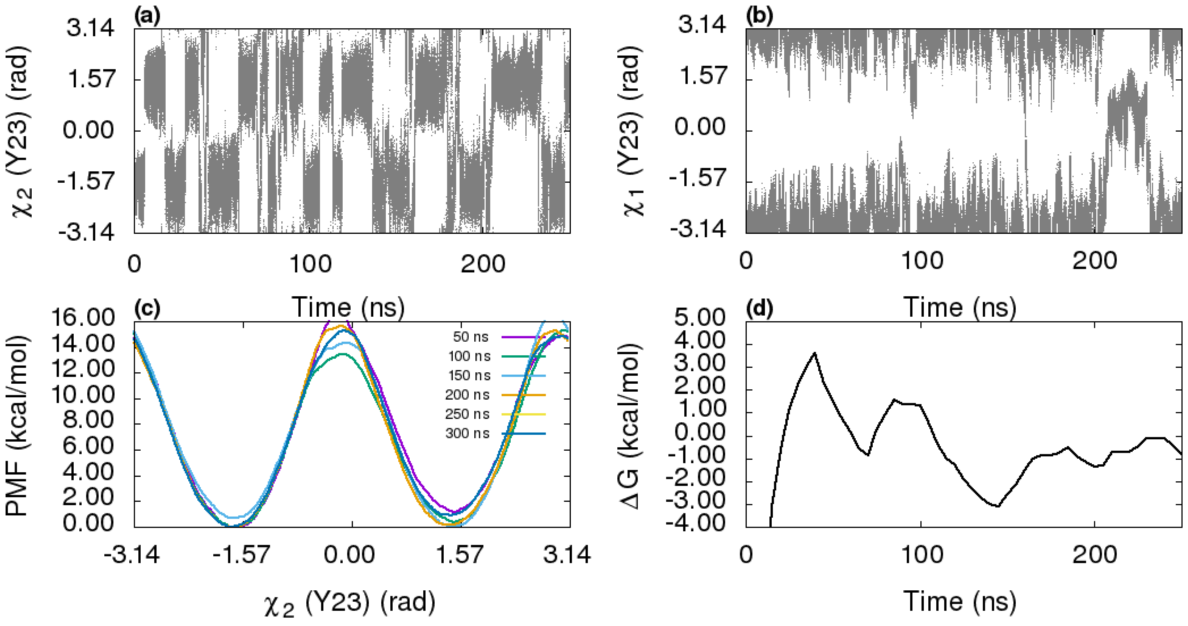


**Figure S32.** The time series of $\chi_2$ and $\chi_1$ during $\chi_2 - \chi_1$metadynamics simulation of residue Y23 in figures (a) and (b), respectively. (c) the projection of free energy on $\chi_2$ in time intervals of 50 ns. (d) The free energy difference between two symmetric states along $\chi_2$.

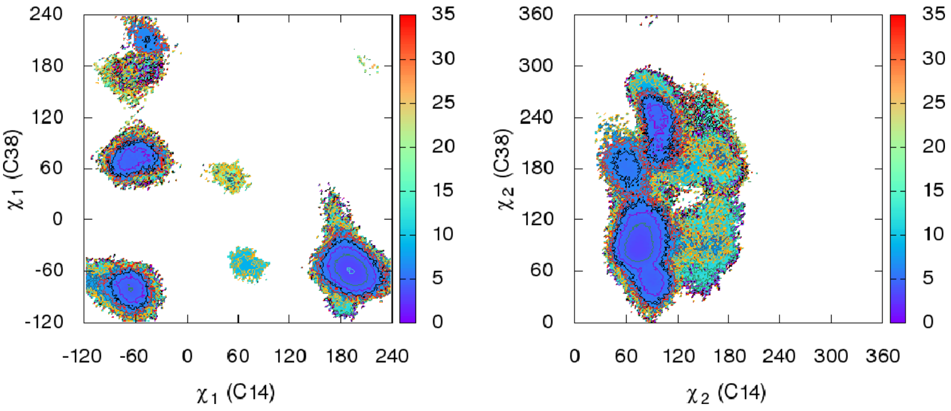

**Figure S33.** The distribution of side-chain torsions of residues C14 and C38 (disulfide bridge C14-C38) in case of non-converged $\chi_2 - \chi_1$metadynamics simulation of residue Y35. The free energy in kcal/mol.

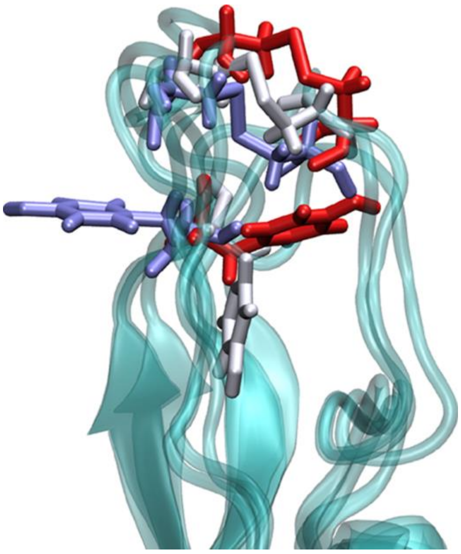

**Figure S34.** The conformations of residue Y35 and C14-C38 disulfide bridge in the native state (red) and unusual conformations (light and dark blue) during well-tempered metadynamics simulations with $\chi_2$, $\chi_1$ of Y35, and $\chi_1$ dihedral of a C14-C38 disulfide bridge.

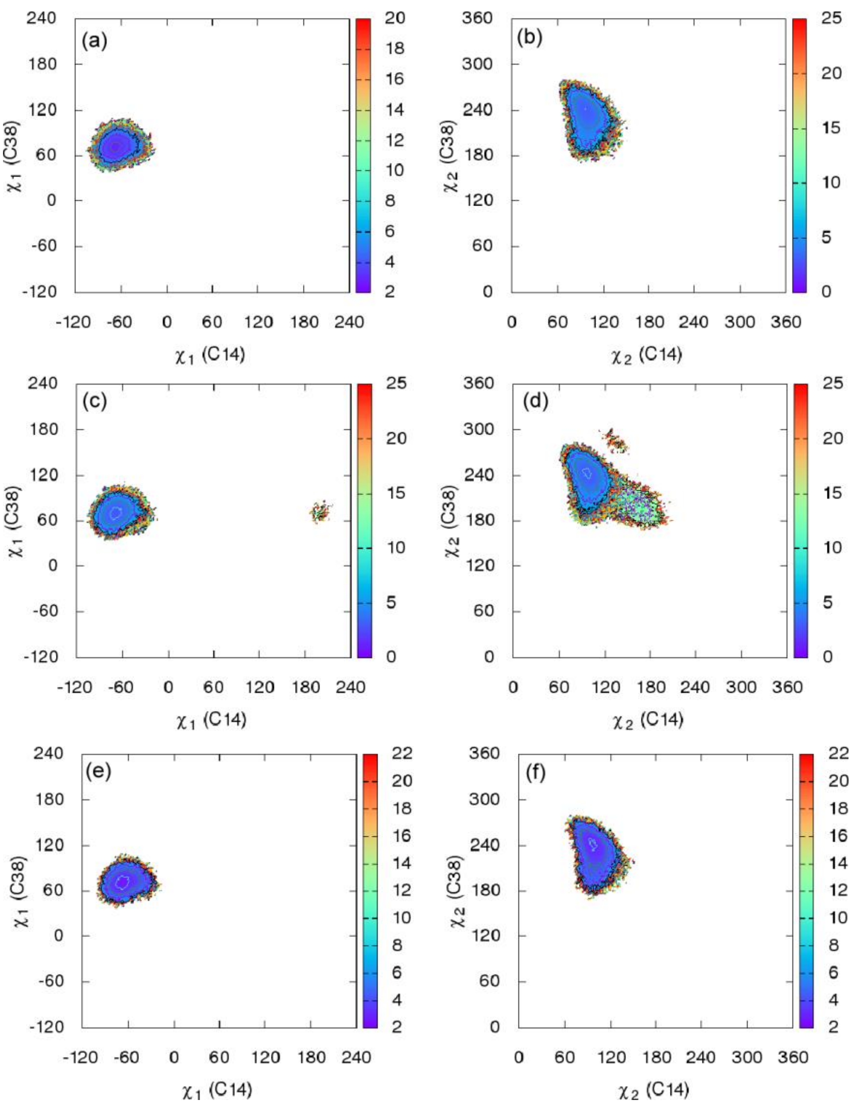


**Figure S35.** The distribution of side-chain torsions of residues C14 and C38 (disulfide bridge C14-C38) in case of each converged metadynamics simulations of fast flipping residues: (a) and (b) residue F4; (c) and (d) Residue Y10; (e) and (f) residue F33. The free energy in kcal/mol.

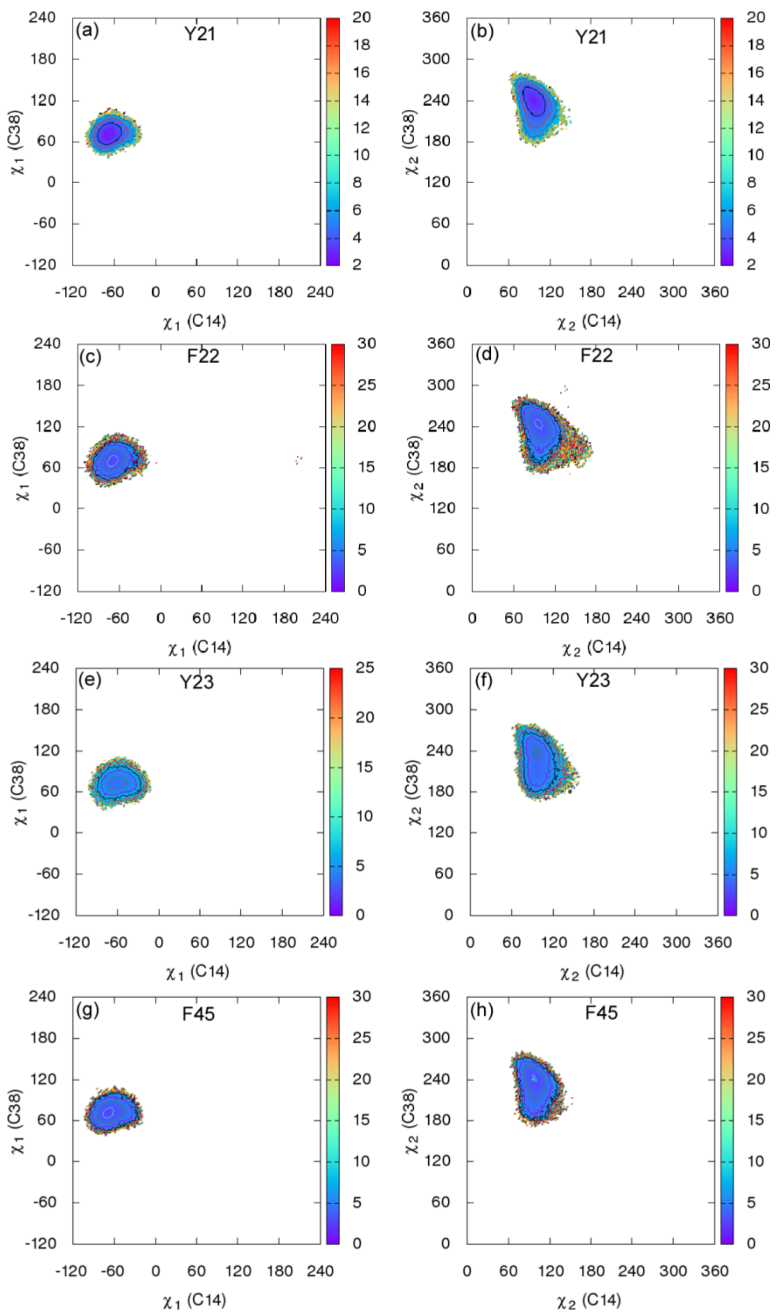


**Figure S36.** The distribution of side-chain torsions of residues C14 and C38 (disulfide bridge C14-C38) in case of each converged metadynamics simulations for residue Y21 (figures (a) and (b); residue F22 (figures (c) and (d)); residue Y23 (figures (e) and (f)); and residue F45 (figures (g) and (h)). The free energy in kcal/mol.

### Solvent Accessible Surface Area (SASA)

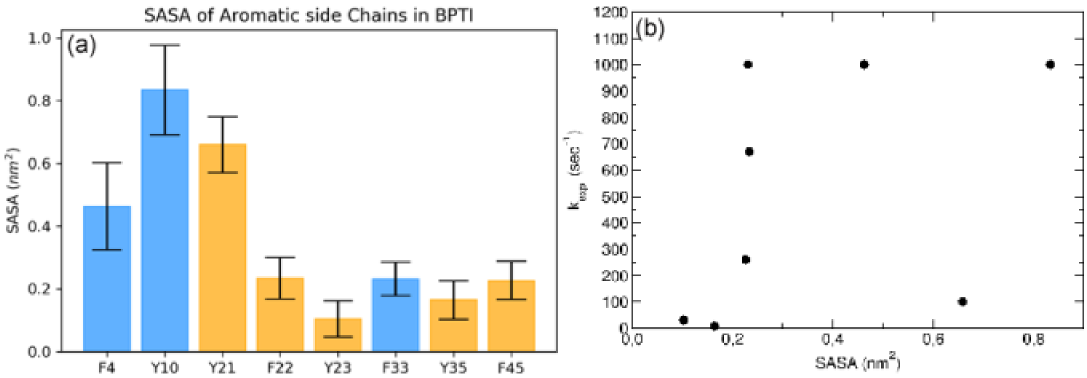


**Figure S37.** (a) SASA for aromatic side chains from standard 100 ns MD simulation. The slow (orange) and fast (blue) flipping residues are shown in a different color. (b) Correlation between flipping rate ($k_{flip}$) and SASA (R=0.39). The SASA of all eight residues using the *gmx sasa* command in *GROMACS* with a standard solvent probe radius of 1.4 Å.

### Circular Correlation of Side-chain torsions

The circular correlation coefficient[1] for dihedrals is calculated using *astropy* python package[2].

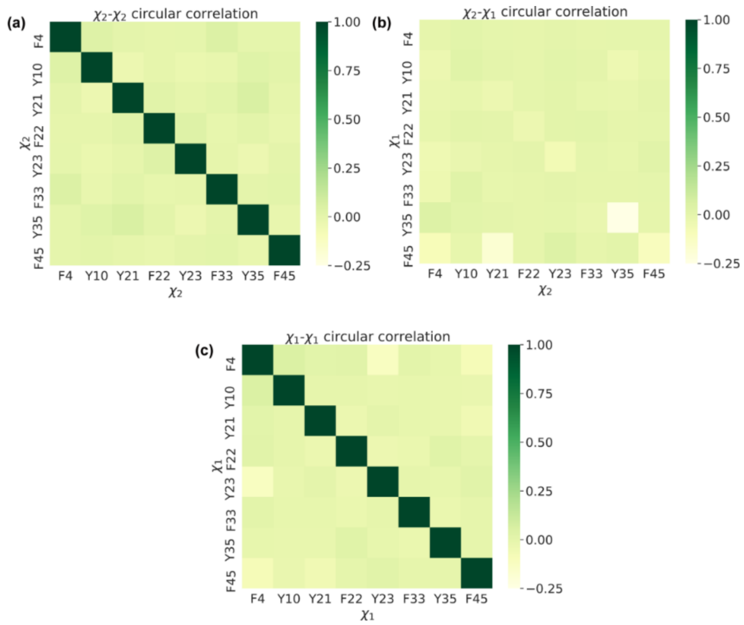


**Figure S38.** The circular correlation coefficient values of torsions $\chi_2$ and $\chi_1$[1] of aromatic residues of BPTI obtained from 1-ms-long trajectory.

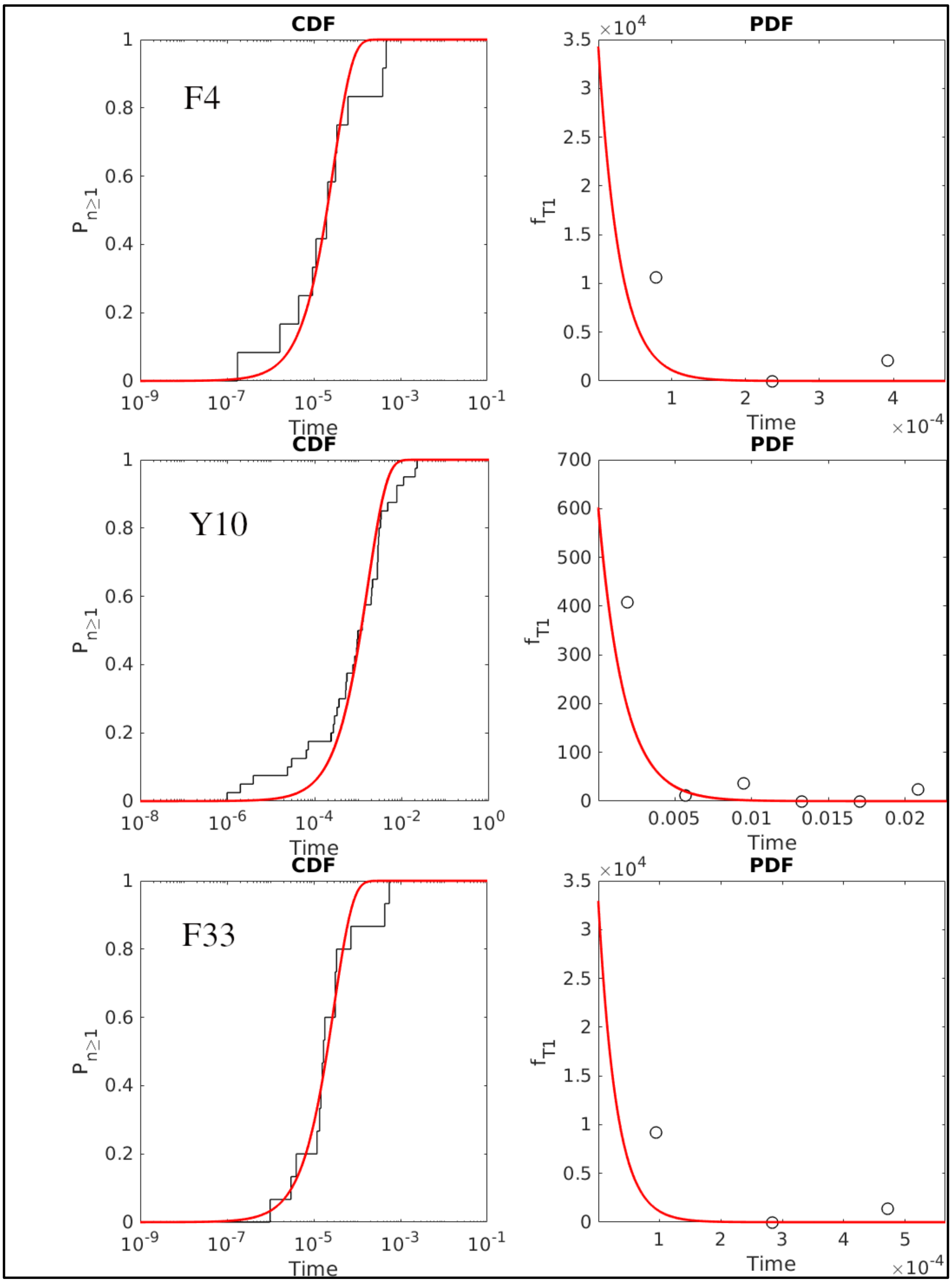


**Figure S39.** Theoretical Poisson fit to the cumulative distribution of transition timescales obtained **from $\chi_2$ infrequent metadynamics** simulations of residue F4, Y10 and F33.

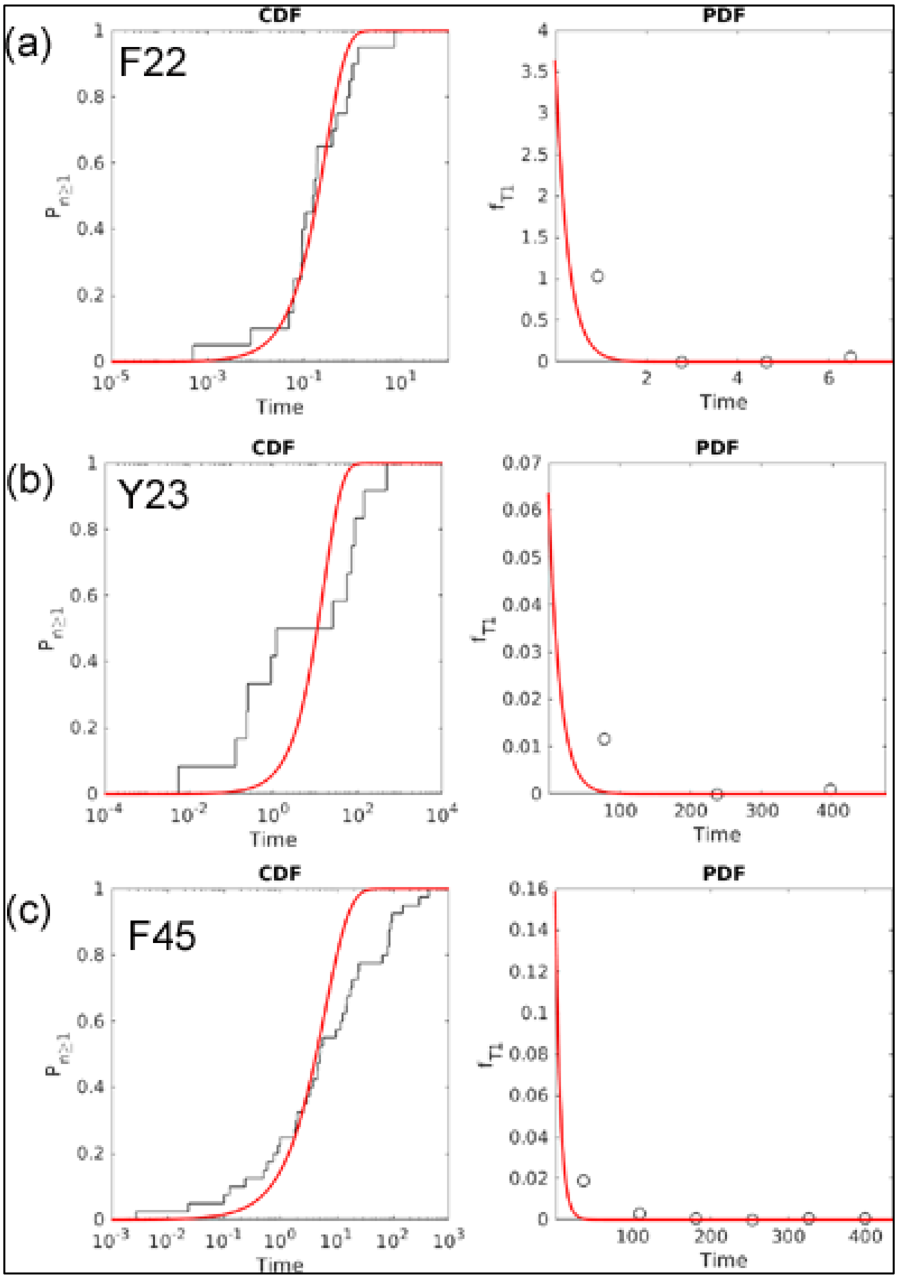


**Figure S40.** Theoretical Poisson fit to the cumulative distribution of transition timescales obtained **from $\chi_2$ infrequent metadynamics** simulations of residue F22, Y23, and F45.

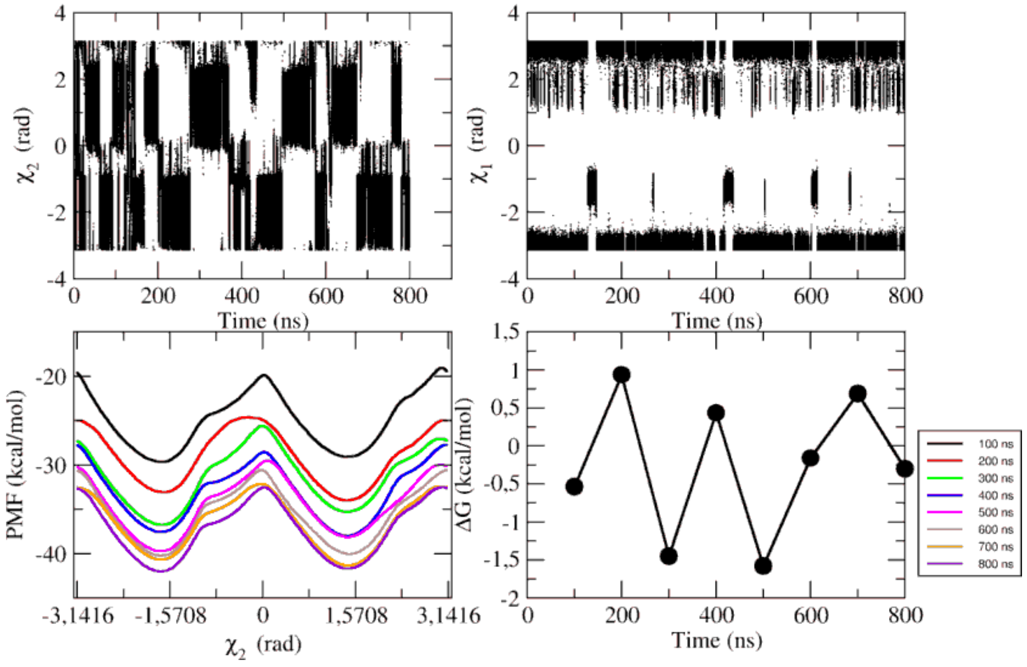


**Figure S41.** Results of $\chi_2$ metadynamics of residue Y10: (a) $\chi_2$ vs. simulation time, (b) $\chi_1$ vs. simulation time, and (c) test of convergence along $\chi_2$ and (d) free energy difference between two identical states. The free energy difference between these two states should be ideally zero as they are identical states.

**CHARMM simulations result:**

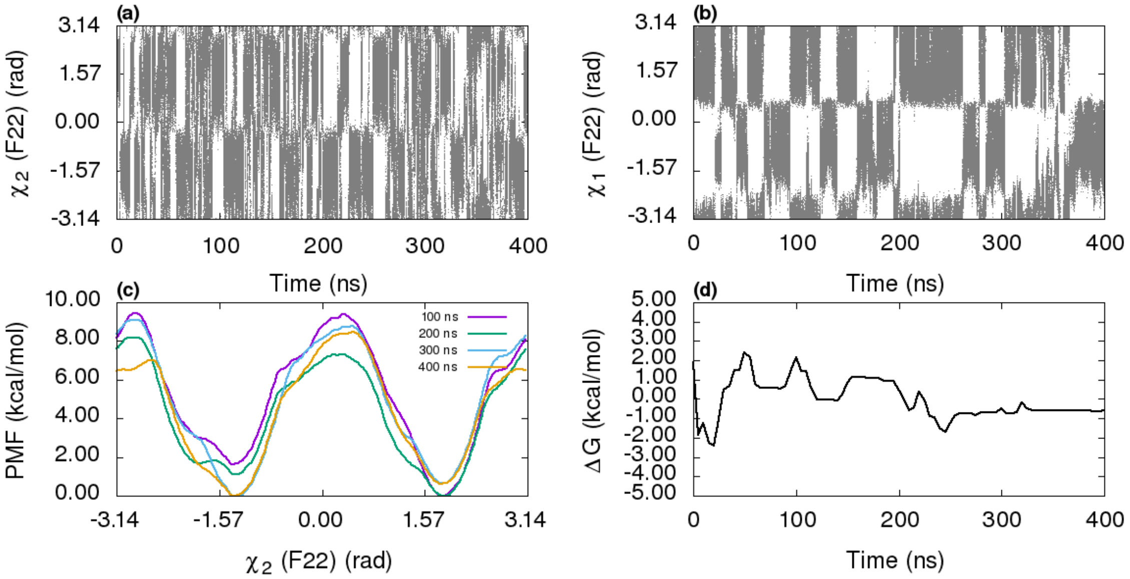


**Figure S42.** Diffusion of $\chi_2$ and $\chi_1$ during $\chi_2 - \chi_1$ metadynamics simulation of residue F22 (with **CHARMM36/standard TIP3P water model**) is shown in figures (a) and (b), respectively. (c) Projection of free energy on $\chi_2$ in time intervals of 100 ns. (d) The free energy difference between two symmetric states along $\chi_2$.

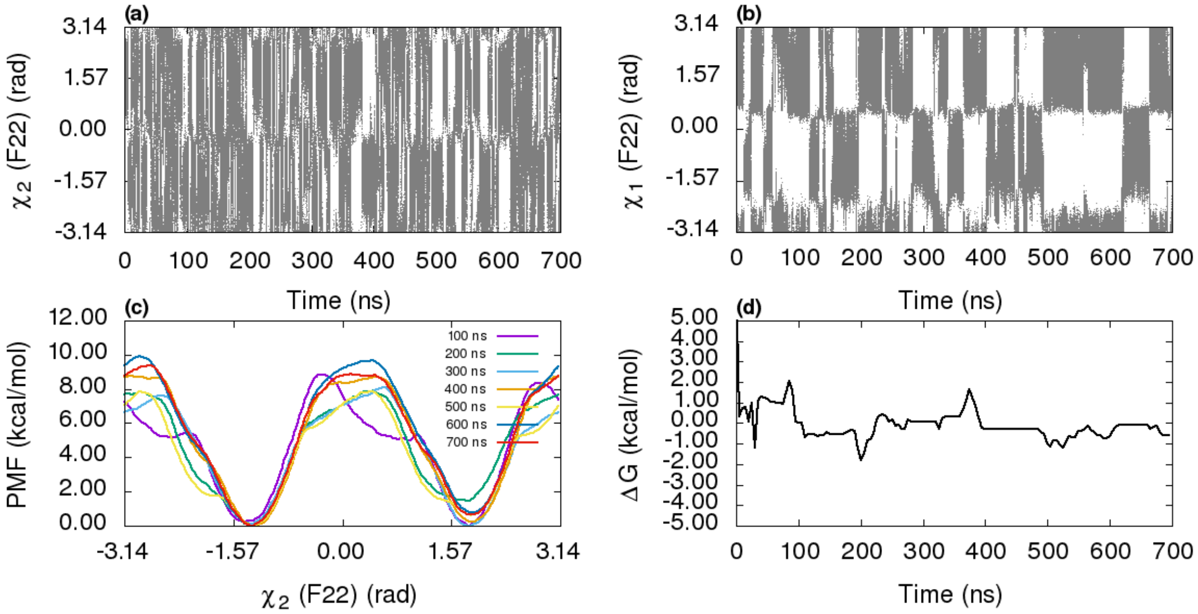


**Figure S43.** Diffusion of $\chi_2$ and $\chi_1$ during $\chi_2 - \chi_1$ metadynamics simulation of residue F22 (with **CHARMM36m/charmm TIP3P water model**) is shown in figures (a) and (b), respectively. (c) Projection of free energy on $\chi_2$ in time intervals of 100 ns. (d) The free energy difference between two symmetric states along $\chi_2$.

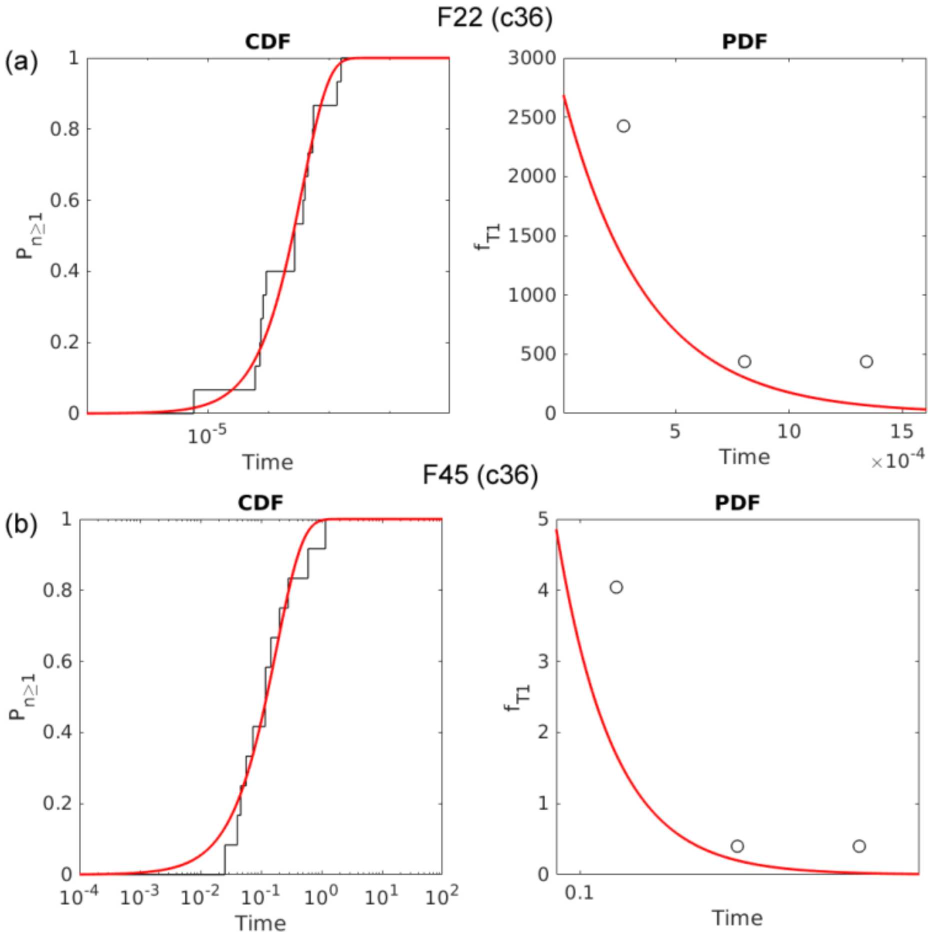


**Figure S44.** The Poisson fit to cumulative distribution function of transition timescales obtained from infrequent metadynamics simulations of CHARMM36/standard TIP3P force field for (a) residue F22 (p-value = 0.69 and $\mu$ ln 2/$t_m$ = 1.24), and (b) residue F45 (p-value = 0.97 and $\mu$ ln 2/$t_m$ = 1.40).

**Appendix I. Residue F22 state-to-state rates.**

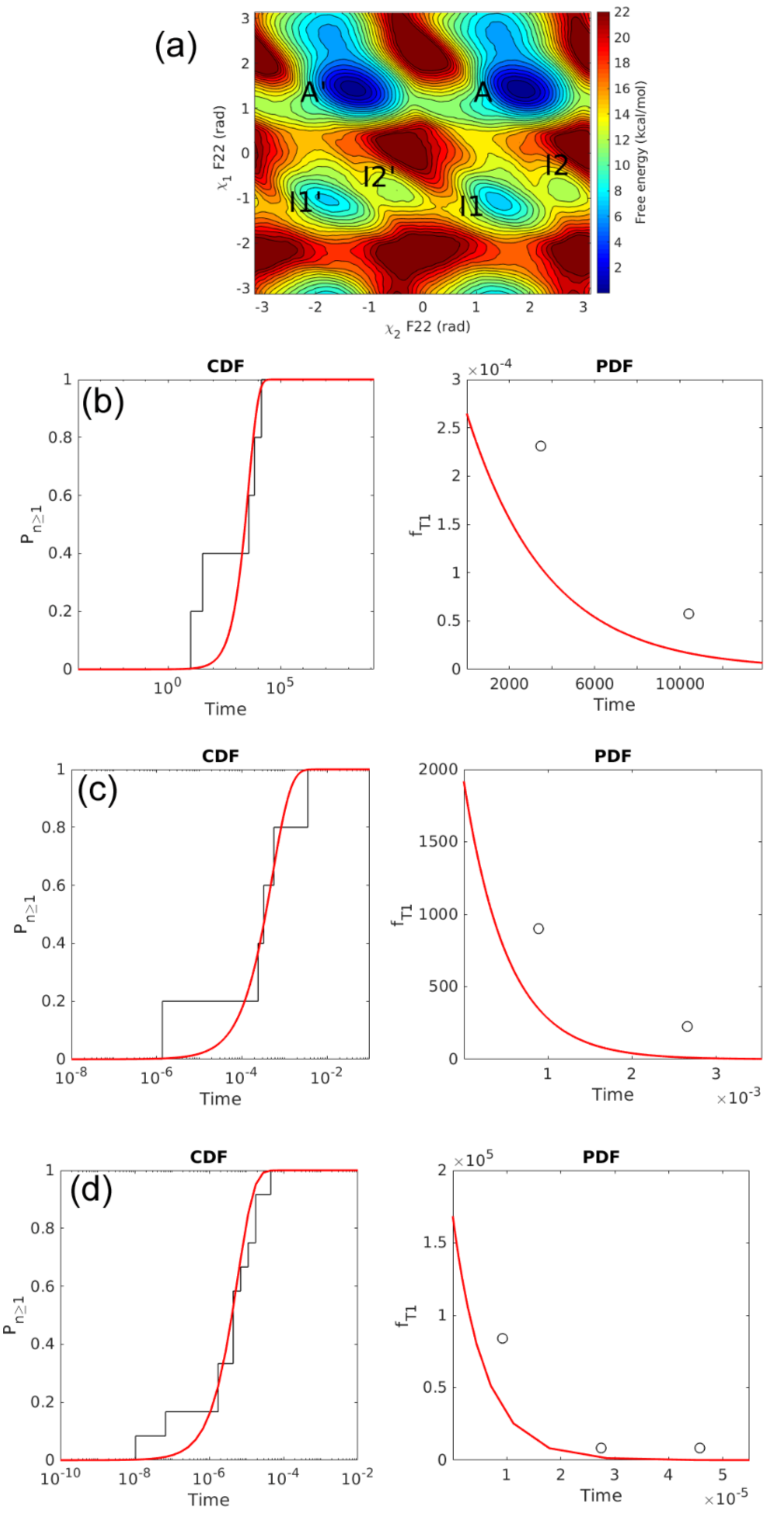


**Figure S45.** The Poisson fit to cumulative distribution function of transition timescales obtained from infrequent metadynamics simulations of (i) A to I1 transitions, (ii) I1 to A transitions, and (iii) I1 to I1' transitions.

Here, we report the results of infrequent metadynamics simulations starting from stable states A, I1 and directed symmetry-equivalent A' and I1' states of residue F22. The trajectories are directed to specific intermediates using harmonica wall bias (see eqn. 3 in main manuscript). The details of wall bias parameters are provided below:

**Table S3.** $k$ and $s_0$ for state-to-state directed InMetaD simulations of residue F22.

| States | $N_{traj}$ | CV | LOWER_WALL ($s_0$) (rad) | UPPER_WALL ($s_0$)(rad) | $k$ (kJ.mol$^{-1}$.rad$^{-2}$) |
|---|---|---|---|---|---|
| A → I1 | 5 | $\chi_2$ | 0.21 | 3.14 | 4184 |
| I1 → A | 5 | $\chi_2$ | 0.21 | 3.14 | 1000 |
| I1 → I1' | 12 | $\chi_1$ | -2.25 | -1.50 | 1000 |

The transitions to nearest neighbors are considered and five simulations were performed in each case. The symmetry of stable state positions on free energy surface allowed to build a complete transition rate matrix with fewer simulations. The infrequent metadynamics simulations are performed for (i) A to I1 transitions, (ii) I1 to A transitions, and (iii) I1 to I1' transitions. The less stable state I2 (and I2') are ignored in the present calculation. The results are reported in Table S4 below.

**Table S4.** Ring flip $t_{flip}(s)$, rate $\mathrm{k}\ (s^{-1})$, p-values, and μ ln 2/tm ratios for state-to-state infrequent metadynamics simulations.

| States | $t_{flip}$ (s) | $\mathrm{k}\ (s^{-1})$ | p-value | μ ln 2/$t_m$ |
|---|---|---|---|---|
| A → I1 | $3.77 \times 10^3$ | $2.65\ \times 10^{-4}$ | 0.33 | 0.89 |
| I1 → A | $5.20\ \times 10^{-4}$ | $1.92\ \times 10^3$ | 0.98 | 1.98 |
| I1 → I1' | $5.93\ \times 10^{-6}$ | $1.69\ \times 10^5$ | 0.39 | 1.29 |

The transition rate matrix based on these simulations is reported in Table S5. The diagonal values[3-4] of this matrix are set to negative of the sum of row elements. All eigenvalues are negative real values as suggested in reference 3. The dominant eigenvalue $83.0\ s^{-1}$ was observed.

**Table S5.** State-to-state transition rate matrix (values in $sec^{-1}$) between nearest-neighbor states A, I1, I1', and A' of residue F22 (AMBER ff14SB) simulations.

| Starting state (row) – end state (column) | A | I1 | A' | I1' |
|---|---|---|---|---|
| A | 0 | $2.65 \times 10^{-4}$ | $4.15 \times 10^{1}$ | 0 |
| I1 | $1.92 \times 10^{3}$ | 0 | 0 | $1.69 \times 10^{5}$ |
| A' | $4.15 \times 10^{1}$ | 0 | 0 | $2.65 \times 10^{-4}$ |
| I1' | 0 | $1.69 \times 10^{5}$ | $1.92 \times 10^{3}$ | 0 |

**Appendix II. Details of VAC derived reaction coordinates.**

We observed transient interactions between residues F22, F33, and P9 of BPTI in metadynamics trajectories of $\chi_2 - \chi_1$ biased simulations of both F22 and F33 (ff14SB force field). Thus, assuming these interactions as a possible reason for recrossing events in $\chi_2 - \chi_1$ infrequently biased simulations, VAC-optimized CV based simulations were performed. We have followed the protocol mentioned in reference [5] (plumID:19.046 at https://www.plumed-nest.org/), and the file for independent component analysis is obtained from the GitHub[6] repository https://github.com/helloyesterday/PLUMED2_TICA.The basis set of following descriptors are used to define a linear combination.

**Table S6.** List of descriptors.

| Residue | Descriptors |
|---|---|
| F22 | $\alpha\beta(\chi_2^{F22})$, $\alpha\beta(\chi_1^{F22})$, $\alpha\beta(\chi_1^{F33})$, $d_1$ |
| F33 | $\alpha\beta(\chi_2^{F33})$, $\alpha\beta(\chi_1^{F33})$, $\alpha\beta(\chi_2^{F22})$, $d_2$ |

$\alpha\beta(\theta)$ is a function transformation of periodic dihedral $\theta$ (in radians) defined as

$$\alpha\beta(\theta) = \frac{1}{2}[1 + \cos(\theta - \theta^{ref})]$$

$\theta^{ref}$= 1.2 rad was used following work in Ref. 7. $\chi_2$ and $\chi_1$ side-chain torsions of residues F22 and F33 are denoted with superscript residue names ($\boldsymbol{\chi_2^{F22}}$, $\boldsymbol{\chi_1^{F22}}$, $\boldsymbol{\chi_2^{F33}}$, $\boldsymbol{\chi_1^{F33}}$). $d_1$ and $d_2$ are distances between atoms of residues P9 and F33.

**$d_1$** = distance between CG atom of P9 and CB atom of F33.

**$d_2$** = distance between C atom of P9 and CB atom of F33.

We used the first 200 ns (F22) and 70 ns (F33) of $\chi_2$ - $\chi_1$ metadynamics simulations based on quasi-equilibrium region, realized by c(t) value, as shown in **figure S10**. The decay of eigenvalues ($\lambda_i, i = 1,2,3,4$ due to 4 CVs) is shown figure below and based on the spectral gap ($\lambda_i - \lambda_{i+1}$) between consecutive eigenvalues.

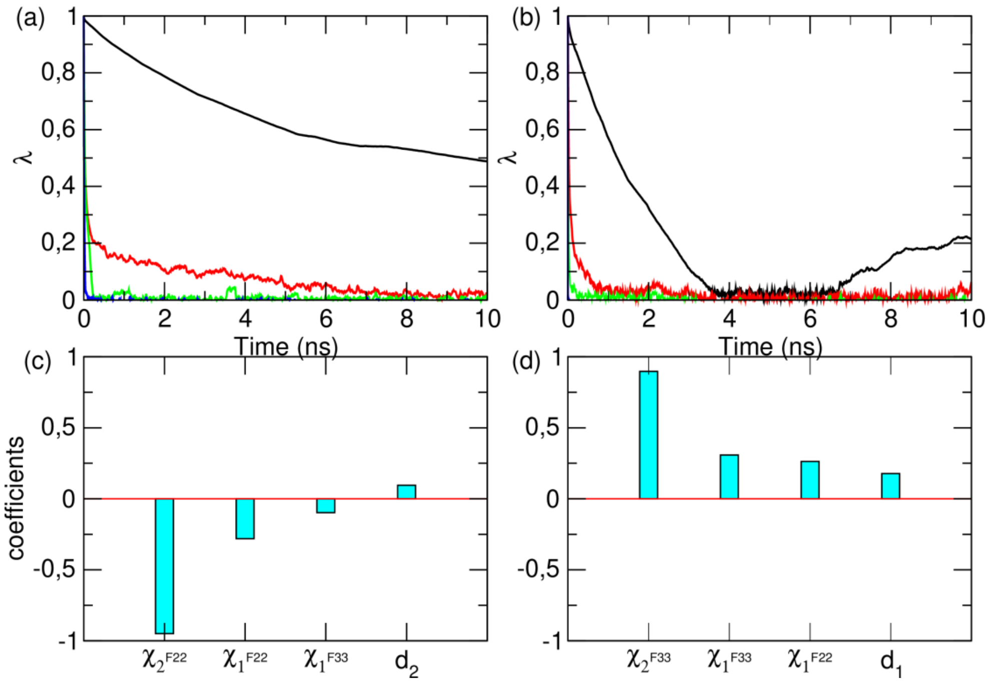


**Figure S46.** The decay of eigenvalues with respect to lag time values for F22 and F33 (figures (a) and (b), respectively). The coefficients of descriptors defining eigenvectors $S_1^{F22}$ and $S_1^{F33}$ (figures (c) and (d), respectively). The coefficients were chosen based on the maximum spectral gap value between the first (black line) and second (red) eigenvectors. The third and fourth eigenvectors are shown with a green and blue line.

The optimal CV ($S_1^{F22}$ and $S_1^{F33}$) used to perform infrequent metadynamics analysis are defined as $S_1^{F22} = -0.95\alpha\beta(\chi_2^{F22}) - 0.28\alpha\beta(\chi_1^{F22}) - 0.097\alpha\beta(\chi_1^{F33}) - 0.095d_2$ and $S_1^{F33} = 0.89\alpha\beta(\chi_2^{F33}) + 0.31\alpha\beta(\chi_1^{F33}) + 0.26\alpha\beta + 0.18d_1$. The cumulative distribution functions of transition timescales obtained by infrequently biasing optimal CVs are shown in **figure S47** below.

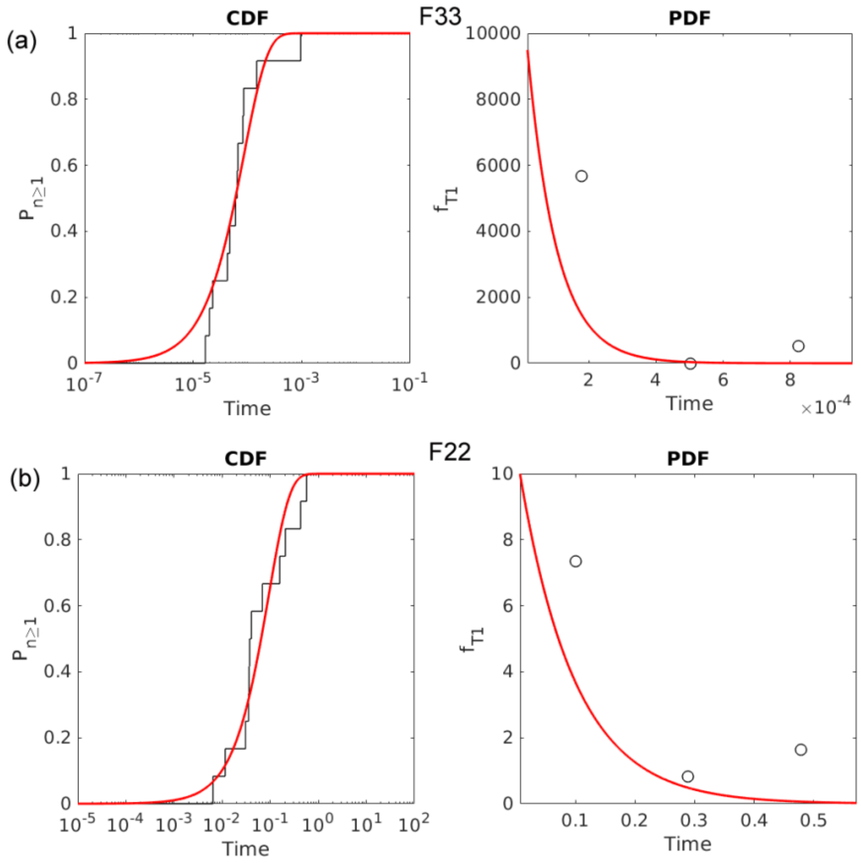


**Figure S47.** The cumulative distribution function of transition timescales obtained from VAC infrequent metadynamics simulations for (a) residue F33 (p-value = 0.70 and $\mu$ ln $2/t_m$ = 1.50), and (b) residue F22 (p-value = 0.49 and $\mu$ ln $2/t_m$ = 2.42).
